# How aging reshapes implicit learning: behavioral and oscillatory evidence from an EEG-adapted Fast Mapping paradigm

**DOI:** 10.64898/2026.10.07.757312

**Authors:** Florentine Fricker, Vincent Dornier, Giulia Buzi, Maelys Morantin, Franck Doidy, Patrice Clochon, Annabelle Goujon, Emmanuel J. Barbeau, Thomas Hinault

**Affiliations:** Université de Caen Normandie, Inserm, EPHE-PSL, PSL Université Paris, CHU de Caen, GIP Cyceron, U1077, NIMH, 14000 Caen, France; Centre de Recherche Cerveau et Cognition (Cerco), UMR5549, Université de Toulouse, 31000 Toulouse, France; Laboratoire de Recherches Intégratives en Neurosciences et Psychologie Cognitive, UMR 1322, INSERM Université de Franche-Comté, 25000 Besançon, France

**Keywords:** Aging, Implicit Learning, Memory, EEG, Theta oscillations, Fast Mapping

## Abstract

Most of everyday memory formation occurs without deliberate intent, yet the neural mechanisms supporting implicit encoding and their reorganization with age remain poorly understood. The present study investigated age-related changes in implicit learning using an EEG-adapted Fast Mapping paradigm (the Disjunctive Inference Implicit Learning Test, DIILT) in 31 younger and 35 older adults, paired with a comprehensive neuropsychological assessment and a one-month delayed recognition measure. Behaviorally, younger adults rapidly acquired novel implicit associations across the learning phase, whereas older adults showed persistent reliance on familiar items and limited improvement for novel associations. Critically, both groups successfully retained novel associations after one month, with implicitly learned associations showing greater long-term stability than explicitly encoded ones regardless of age. At the neural level, younger adults exhibited early posterior theta engagement for novel compared to familiar items, which disappeared by the end of learning, consistent with novelty-driven associative encoding. Older adults showed no such theta modulation for novel items, pointing to reorganization of the neural mechanisms supporting implicit encoding. Theta dynamics predicted immediate recognition in younger adults but not in older adults who rather showed theta association with processing speed, suggesting an age-related shift from memory-specific binding to domain-general cognitive resource allocation, consistent with neural maintenance frameworks of cognitive aging. These findings demonstrate that implicit encoding produces durable memory traces across the lifespan but relies on reorganized oscillatory dynamics in aging, with implications for understanding how cognitive aging reshapes everyday memory formation.

## 1. Introduction

In everyday life, we rarely intend to memorize new information explicitly. Instead, we rather incidentally extract regularities, bind events to their context, and only later reconstruct what we learned and use it specifically (Goujon et al., 2015; Theeuwes et al., 2022; Wagnon et al., 2019; Xue, 2022; Yonelinas et al., 2019). These implicit encoding processes are central to how we make sense of experience. Yet despite their relevance to understand everyday cognition, most research on memory in aging has focused on intentional encoding and explicit retrieval, leaving open how aging reshapes these largely implicit learning mechanisms that structure experience outside the laboratory. Addressing this issue is important not only for theoretical advances, but also for characterizing cognitive aging through paradigms that tap into encoding processes closer to those operating in naturalistic settings. Yet studying these processes at the neural level remains challenging, as capturing their oscillatory signature requires repeated exposures to the same associations, a constraint rarely met by paradigms designed to evaluate implicit learning. To fill this gap, the present study investigated the age-related differences in implicit encoding using an EEG-adapted Fast Mapping paradigm, paired with a comprehensive neuropsychological assessment, in younger and older adults. This design enabled the simultaneous evaluation of behavioral performance and neural oscillatory dynamics during acquisition of novel information, providing a unique opportunity to characterize how neural mechanisms supporting implicit learning evolve with advancing age.

Cognitive aging is marked by profound inter-individual variability and a differential gradient of cognitive decline (see Nyberg et al., 2020, for a review; Habib et al., 2007; Lindenberger, 2006; Salthouse, 2010). While episodic and working memory typically show the steepest age-related decline (Gorbach et al., 2017), retrieval in semantic memory remains comparatively stable with advancing age (Hedden & Gabrieli, 2004; Nyberg et al., 2020). Moreover, compensatory neural recruitment frequently emerges alongside maintenance of memory performance in some individuals (Jauny et al., 2022; Park & Reuter-Lorenz, 2009). This dissociation suggests that aging does not uniformly impair memory but rather alters the mechanisms supporting new memories formation. Yet how aging affects implicit learning processes that structure everyday experience, and more specifically reshapes the oscillatory dynamics underlying implicit encoding of novel associations remains largely uncharacterized. A central hypothesis of the present study is that certain aspects of semantic learning are supported by statistical learning mechanisms. Statistical learning provides a framework for examining how one extracts environmental regularities without explicit instruction (Frost et al., 2015; Saffran et al., 1996; see Aslin & Saffran, 2025, for a review) and refers to the ability to spontaneously detect transitional probabilities and infer latent structure from continuous sensory input across space and time. Using an adapted Fast Mapping procedure, the goal of the present study was to extend this logic to semantic knowledge acquisition. This exclusion-based learning procedure relies on novel labels being inferred from their co-occurrence with known referents by disjunctive syllogism mechanism (Sharon et al., 2011). Studies have proposed that Fast Mapping may operationalize implicit semantic binding, whereby new associations are integrated through contextual inference rather than deliberate memorization (Cooper et al., 2019a; Merhav et al., 2015; Vasilyeva et al., 2019). Of note, memory for fast-mapped associations has typically been assessed over short delays, as is common in standard memory assessments, raising the question of whether such implicitly formed traces persist over longer periods of time that are more representative of everyday memory.

Over the past decade, the cognitive mechanisms underlying Fast Mapping have begun to be characterized more precisely. Beyond the exclusion-based inference mechanism itself, successful Fast Mapping appears to draw on several interacting cognitive processes. The rapid integration of novel associations into existing semantic networks suggests that prior knowledge plays a key facilitating role: new labels are anchored to pre-existing conceptual representations, allowing efficient encoding without deliberate memorization (Coutanche & Thompson-Schill, 2014; Merhav et al., 2015). Notably, the degree of semantic overlap between novel and familiar items has been shown to modulate the extent to which this rapid cortical integration can occur (Zaiser et al., 2022). Additionally, Fast Mapping requires selective attention toward novelty (Constant et al., 2023). The learner must direct attentional resources toward the unfamiliar items while inhibiting the known distractor, a process that places demands on attentional and executive resources (Diamond, 2013; Miyake et al., 2000). These cognitive demands suggest that Fast Mapping, while incidental in nature, is not purely automatic and that its efficiency depends on the availability of attentional, executive, and mnemonic resources at encoding. As aging is known to differentially impact these cognitive systems, these aspects have direct implications for how Fast Mapping may be expressed in aging.

The neural mechanisms underlying this implicit encoding process, especially in older population, remain incompletely understood. Electroencephalography (EEG), with its millisecond temporal resolution, is particularly well-suited to track neural dynamics as they unfold during learning (Hinault et al., 2021; Merkin et al., 2023; Perinelli et al., 2022) and could help in shedding light on these implicit encoding processes. Theta band dynamics, in particular, represent a plausible electrophysiological substrate, as they have been consistently implicated in novelty-driven associative encoding and successful memory formation (Backus et al., 2016; Biba et al., 2026; Herweg et al., 2020; Lega et al., 2012). In that matter, recent EEG work has directly implicated theta in the neural indexing of novel word-meaning associations before and after a Fast Mapping exposure (Shtyrov, 2011; Shtyrov et al., 2021, 2022; Vasilyeva et al., 2019), suggesting that even a single implicit learning episode leaves measurable traces in cortical activity. Growing interest in the electrophysiological correlates of Fast Mapping has recently extended to retrieval processes, with event-related potential results indicating that associations learned through Fast Mapping are later retrieved through familiarity-based mechanisms distinct from those supporting explicit encoding (Festag et al., 2026). However, across this body of work, EEG has so far been used to characterize neural responses before, after, or during retrieval of Fast Mapping associations, rather than oscillatory dynamics unfolding during the encoding episode itself. Consequently, to our knowledge, no study has yet recorded time-frequency dynamics in real time during Fast Mapping learning, leaving the real-time theta substrate of implicit encoding largely uncharacterized. Moreover, given the well-documented shifts in oscillatory dynamics that accompany healthy aging (Hinault et al., 2021), an unequally understudied question is whether the neural mechanisms supporting implicit encoding are preserved or reorganized with age, potentially extending to frequency bands beyond theta.

Previous literature suggests that implicit learning remains functionally available in aging, but that aging could change which systems support this process and could alter stability and expression of implicitly acquired information, especially under high attentional or strategic demands (Dennis & Cabeza, 2011; Rieckmann & Bäckman, 2009; Schevenels et al., 2023; Wagnon et al., 2019). However, these effects in older adults are inconsistent, raising the possibility that aging alters the neural implementation of implicit encoding rather than abolishing it. Moreover, direct comparisons between younger and older adults during Fast Mapping encoding remain scarce, limiting our understanding of how age modulates the mechanisms involved during learning itself rather than at retrieval, and how theta-dependent encoding mechanisms are expressed, and potentially reorganized, across age groups.

Taken together, these findings converge on a central question: how does aging reshape the oscillatory mechanisms supporting implicit encoding in a Fast Mapping procedure, and what are the behavioral and cognitive consequences of this reorganization? To shed light on this question, the present study was designed to address these two primary objectives. First, we characterized implicit learning across age groups using an EEG-adapted Fast Mapping paradigm (called the Disjunctive Inference Implicit Learning Test, or DIILT, as this paradigm departs from the classing Fast Mapping task in order to comply with EEG constraints), focusing on behavioral performance and task-related theta dynamics over the course of learning. Second, we examined the relationships between task-related oscillatory dynamics, subsequent memory performance, and individual neuropsychological scores, probing whether these associations differ across age groups. We hypothesized that younger adults would exhibit increased theta activity for novel associations that would decrease as learning stabilizes, consistent with novelty-driven encoding. Older adults, however, were expected to show attenuated or redistributed theta modulation, consistent with the oscillatory slowing characteristic of healthy aging, or compensatory recruitment/maintenance (Jauny et al., 2022). At the behavioral level, we assumed that both groups would successfully encode novel associations implicitly, but that older adults would show more difficulty acquiring these associations throughout learning. Finally, we expected that theta dynamics during encoding would be related to subsequent memory performance, with the nature of this association differing between age groups, reflecting a reorganization of the neural substrates supporting implicit learning with age.

By examining implicit learning and its oscillatory substrates within a lifespan framework, this study aimed to clarify how the brain rapidly and implicitly forms new memory traces, and how this process evolves with age.

## 2. Method

### 2.1. Participants

Seventy-one French-speaking volunteers participated in the experiment. All participants reported normal or corrected-to-normal vision and hearing, and no history of major medical conditions, neurological or psychiatric disorders, or head trauma. Older adults were screened using the Montreal Cognitive Assessment ( MoCA; Nasreddine et al., 2005) and were included only if they scored ≥ 26 (mean score = 28.38 ± 1.16).

The study was approved by a French ethics committee (Comité de Protection des Personnes, CPP; ID-RCB: 2024-A00167-40), and all participants provided oral and written informed consent in accordance with the Declaration of Helsinki (2024). Participants were recruited through academic mailing lists and social media and received 20 € for their participation.

Five participants were excluded from the final analyses: three due to excessive EEG noise, one because the participant misunderstood the learning-phase instructions of the implicit learning task, and one due to a technical issue during acquisition.

The final sample consisted of 66 participants: 31 younger adults (16 females; mean age = 23.94 ± 2.89 years, ranging from 20 to 31 years old) and 35 older adults (26 females; mean age = 69.26 ± 5.54 years, ranging from 61 to 79 years old).

Furthermore, two participants (one younger adult and one older adult) did not complete the second session and were consequently excluded from analyses involving delayed recognition but were retained for first-session analyses.

### 2.2. Procedure

Participants completed two experimental sessions separated by approximately one month (mean interval = 30.45 days, min = 18, max = 61). Each session lasted approximately three hours. All behavioral tasks were run on Opensesame (version 4.1.7; Mathôt et al., 2012).

#### Session 1

During the first session, older adults were screened using the Montreal Cognitive Assessment (MoCA) to confirm eligibility. Before the implicit learning task (DIILT), the researcher gave instructions to the participants, then completed a training block composed of four trials. The training was repeated if requested. Data from the training session were not included in the analyses.

After EEG installation, participants were seated approximately 90 cm from a computer screen inside a faraday cage and performed the implicit learning task under EEG recording, followed by the immediate recognition task, which was not EEG recorded due to time constraints.

After EEG removal, participants completed the explicit encoding learning task and its immediate recognition on a computer in a quiet testing room. Finally, they completed the first part of the neuropsychological assessment battery.

#### Session 2

During the second session, participants first completed the one-month delayed memory recognition tasks related to both the implicit and explicit learning tasks, presented randomly across participants. Then they completed the remaining part of the neuropsychological assessment battery. The overall experimental timeline is illustrated in Figure 1.

**Figure 1.**
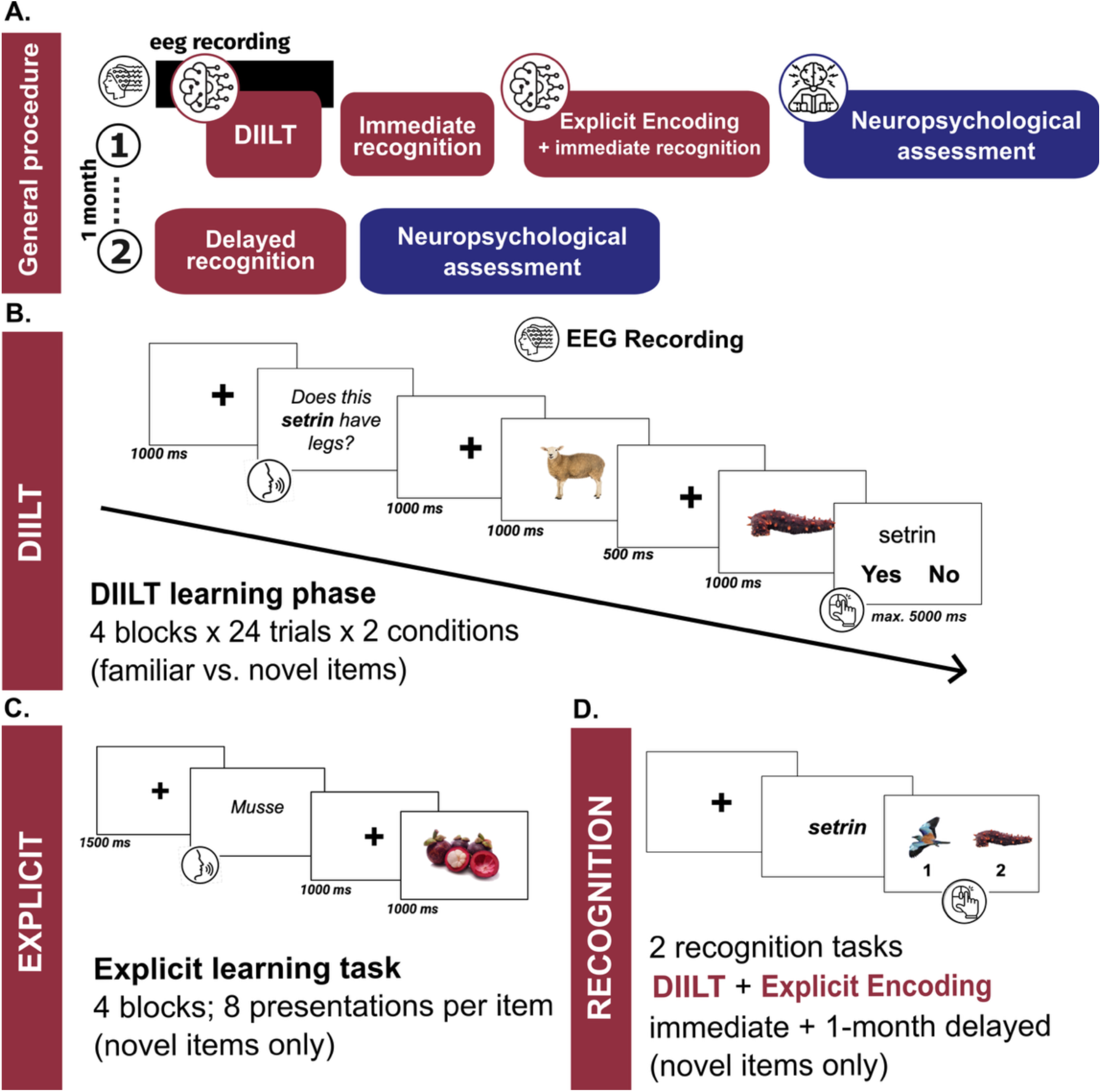
General experimental procedure sessions and overview of the DIILT and the explicit experimental paradigms. (A) Participants completed two sessions separated by approximately one month. Session 1 comprised the implicit learning phase (DIILT) under EEG recording, and its immediate recognition task (not EEG recorded). After EEG removal, they performed the explicit encoding task and its immediate recognition task after removal and the first part of the neuropsychological battery. Session 2 included the one-month delayed recognition tasks for both the implicit and explicit tasks and completion of the remaining neuropsychological assessments. (B) *Implicit encoding phase*. Participants completed four blocks of 24 trials under EEG recording, including two conditions: familiar associations and novel associations. On each trial, a spoken question referred to one of two sequentially presented visual stimuli. Participants answered the yes/no question using the mouse. (C) *Explicit encoding phase.* Participants completed four blocks comprising twelve novel items only, each presented twice per block for a total of eight presentations. Each trial consisted of the auditory presentation of a pseudoword followed by its corresponding image. Participants were explicitly instructed to memorize the image-word associations and knew they would have to recall them after the task. (D) *Recognition.* Memory was assessed twice (after a 5-min delay post learning and again one month later during the second session) using a two-alternative forced-choice task for novel items learned in both the implicit and explicit tasks. Each trial began with the simultaneous auditory and visual presentation of a pseudoword, followed by two previously learned images displayed side by side. Participants were instructed to select the image corresponding to the novel word using the mouse.

### 2.3. Experimental tasks

#### 2.3.1. Implicit encoding of novel information: the Disjunctive Inference Implicit Learning Test (DIILT)

##### 2.3.1.1. DIILT: Learning phase

To assess implicit encoding, we created a new paradigm adapted from classical Fast Mapping tasks to counteract EEG constraints: the Disjunctive Inference Implicit Learning Test (DIILT; see Figure 1B). The DIILT was presented as a perceptual task lasting approximately 30-min under EEG recording. Note that participants always started with the implicit procedure to minimize memory strategy during learning. It comprised four blocks of 48 trials, including two conditions: familiar and novel associations (see Supplementary Table S1 for full item list and details).

On each trial, a spoken question referring to perceptual characteristics (e.g., appearance, material, behavior, etc.) was played through a speaker (e.g., familiar: “Is this *sheep* black?”; novel: “Does this *setrin* have legs?”). Two visual stimuli were then displayed sequentially in a random order for 1000 ms each, followed by a response screen. The question always referred to one of the two images and required a yes or no response. Participants answered using the mouse with their right hand (left click = yes; right click = no). The response screen remained on display until an answer was provided, up to a maximum of 5000 ms, before the next trial began.

The pair of visual stimuli consisted of a target (the image corresponding to the spoken question) and a referent (a familiar image unrelated to the question), separated by a 500 ms fixation cross. For example, in a novel item trial with the question “Does this Crute have a tail?”, followed by an image of a strawberry and an unfamiliar object, the strawberry serves as the referent and the unfamiliar object as the target. Participants would thus have to infer by disjunctive syllogism that the unfamiliar object corresponds to the *Crute*. Between each block participant were allowed short breaks before manually engaging in the questions again when ready (generally taking an average of 30 seconds to a minute).

###### Question selection

Eight questions were created per item (two per block) using synthetic speech (TTSMaker; https://ttsmaker.com/; Dore’s voice). The structure was consistent across trials: « Does this / Is this » + familiar or novel item name + characteristic. Questions lasted approximately one second on average. A *t*-test on the number of letters per question revealed no difference in length between novel and familiar items (*t*(262.99) = 0.82, *p* =.41, n = 384). Half of questions expected a *yes* answer and the other half a *no* answer.

###### Item selection

Visual stimuli were centrally displayed on a white background (approximately 10 x 10 cm on a 47.6 cm x 26.8 cm monitor). Brightness and sound volume were kept constant across participants.

###### Randomization

The two novel item lists were counterbalanced across implicit and explicit tasks according to participant number. Block order, question order, and distractor assignment were randomized.

Each block comprised 48 trials (24 familiar and 24 novel associations). Each item appeared twice per block with two different questions (one requiring a “yes” response and one requiring a “no” response). In total, participants completed 192 trials.

###### Behavioral data

For each participant, we computed mean accuracy and mean response times per condition and per block.

##### 2.3.1.2. DIILT: Recognition task

After a 5-min delay, participants completed a surprise two-alternative forced-choice recognition test (see Figure 1D) on the 12 novel items learned during DIILT. Each trial began with a 1500 ms fixation cross, followed by the simultaneous visual and auditory presentation of a pseudoword. Two previously learned novel images were then displayed side by side. Participants selected the image corresponding to the pseudoword using the mouse. Images remained on the screen until a response was made, with no time limit, allowing us to collect spontaneous mean response time and mean accuracy for immediate recognition. Memory consolidation was assessed again one month later during the second session using the same recognition procedure.

#### 2.3.2. Explicit encoding

##### 2.3.2.1. Learning phase

Participants completed four blocks twelve novel items (distinct from those used in the DIILT task), each presented twice per block, for a total of eight presentations per item across the four blocks (see Figure 1C). Each trial consisted of the auditory presentation of a pseudoword followed by the corresponding image displayed on a white background for 1000 ms. Visual parameters were identical to the DIILT task. Participants were explicitly instructed to try to memorize the image-word associations, as they would be tested afterward.

##### 2.3.2.2. Recognition task

After a 5-min delay, participant completed the same two-alternative forced-choice recognition task for the explicitly learned novel items. The task was repeated one month later. Mean accuracy and response times were analyzed for both immediate and delayed recognition.

### 2.4. Neuropsychological Assessment and questionnaires

Participants additionally underwent a comprehensive neuropsychological and questionnaire-based assessment to characterize cognitive functioning across domains (see Table 1). Socioeconomic status was assessed with the French adaptation of the Hollingshead’s four-factor Index of Socio-Economic status.

**Table 1.** Descriptive statistics and group comparisons for all neuropsychological and questionnaire measures.

| Tests | n (YA/OA) | Young Adults M (SD) [min, max] | Older Adults M (SD) [min, max] | t(df) | p | Cohen's d |
| --- | --- | --- | --- | --- | --- | --- |
| Age | 31/35 | 23.94 (2.89) [20.00, 31.00] | 69.26 (5.54) [61.00, 79.00] | W=0 | < .001*** | r=-1.00 |
| Gender (F/M) | 31/35 | F: 16 (51.6%), M: 15 (48.4%) | F: 26 (74.3%), M: 9 (25.7%) | $\chi^2(1)=3.65$ | .056 | $\phi=.20$ |
| Socioeconomic Status | 31/35 | 6.91 (1.37) [3.67, 9.00] | 6.65 (1.19) [3.33, 9.33] | t(60)=0.84 | .404 | 0.21 |
| Years of education | 31/35 | 19.32 (1.90) [15.00, 24.00] | 15.37 (3.44) [9.00, 26.00] | W=922 | < .001*** | r=0.70 |
| CVLT free recall (immediate) | 31/35 | 14.52 (1.46) [11.00, 16.00] | 13.74 (2.03) [9.00, 16.00] | W=658 | .131 | r=0.21 |
| CVLT cued (immediate) | 31/35 | 14.65 (1.38) [11.00, 16.00] | 14.09 (1.42) [11.00, 16.00] | W=676 | .078 | r=0.25 |
| CVLT free recall (delayed) | 30/35 | 14.93 (1.11) [12.00, 16.00] | 14.37 (1.73) [10.00, 16.00] | W=602 | .296 | r=0.15 |
| CVLT cued (delayed) | 31/35 | 14.81 (1.49) [11.00, 16.00] | 14.29 (1.67) [11.00, 16.00] | W=650 | .154 | r=0.20 |
| CVLT recognition | 31/34 | 43.42 (0.96) [41.00, 44.00] | 43.29 (1.03) [40.00, 44.00] | W=574 | .478 | r=0.09 |
| **Autobiographical Recollection Test** | 31/35 | 116.61 (18.17) [63.00, 142.00] | 103.37 (17.72) [64.00, 141.00] | t(63)=2.99 | **.004\*\*** | 0.74 |
| **Stroop (reading)** | 31/35 | 1.16 (0.19) [0.90, 1.70] | 1.27 (0.20) [0.95, 1.70] | W=374 | **.031\*** | r=-0.31 |
| **Stroop (interference)** | 30/35 | 1.61 (0.28) [1.23, 2.44] | 2.00 (0.42) [1.21, 2.74] | t(59)=-4.33 | **< .001\*\*\*** | -1.05 |
| **N-Back** | 31/34 | 20.90 (5.35) [8.00, 28.00] | 15.56 (6.17) [5.00, 27.00] | W=786 | **< .001\*\*\*** | r=0.49 |
| **Processing Speed** | 30/35 | 65.00 (11.03) [41.00, 96.00] | 51.89 (11.33) [31.00, 75.00] | t(62)=4.72 | **< .001\*\*\*** | 1.17 |
| **Digit span (forward)** | 31/34 | 5.87 (0.88) [4.00, 8.00] | 5.26 (0.71) [4.00, 7.00] | W=728 | **.005\*\*** | r=0.38 |
| **Digit span (backward)** | 31/35 | 5.71 (1.30) [3.00, 8.00] | 4.77 (1.26) [2.00, 8.00] | W=758 | **.004\*\*** | r=0.40 |
| ASQ total score | 31/34 | 61.61 (11.37) [39.00, 79.00] | 57.24 (9.83) [37.00, 74.00] | t(60)=1.65 | .104 | 0.41 |
| ASQ Internal Orientation | 31/34 | 25.65 (5.91) [14.00, 35.00] | 23.65 (5.29) [14.00, 35.00] | t(60)=1.43 | .158 | 0.36 |
| ASQ External Orientation | 31/34 | 18.03 (6.78) [7.00, 28.00] | 16.21 (4.90) [6.00, 27.00] | W=615 | .250 | r=0.17 |
| BIS-11 total score | 31/34 | 62.87 (10.78) [41.00, 85.00] | 60.21 (7.76) [45.00, 81.00] | t(54)=1.14 | .262 | 0.29 |
| BIS-11 Planning difficulties | 31/34 | 23.13 (4.36) [17.00, 33.00] | 24.41 (4.15) [13.00, 32.00] | W=406 | .114 | r=-0.23 |
| **BIS-11 motor impulsivity** | 31/33 | 21.58 (4.57) [14.00, 31.00] | 19.48 (2.58) [13.00, 25.00] | t(47)=2.24 | **.030\*** | 0.57 |
| **BIS-11 cognitive impulsivity** | 31/34 | 18.16 (4.51) [8.00, 29.00] | 16.00 (2.55) [12.00, 22.00] | t(46)=2.35 | **.023\*** | 0.60 |
| **HADS Anxiety** | 31/34 | 7.74 (3.54) [1.00, 16.00] | 6.03 (2.68) [1.00, 12.00] | t(56)=2.18 | **.033\*** | 0.55 |
| HADS Depression | 30/34 | 3.03 (2.08) [0.00, 7.00] | 3.00 (2.22) [0.00, 9.00] | W=520 | .891 | r=0.02 |
| **GDS** | 31/34 | 7.26 (5.22) [0.00, 20.00] | 3.74 (2.57) [0.00, 11.00] | W=754 | **.003\*\*** | r=0.43 |

**Note.** YA = Younger Adults; OA = Older Adults. CVLT = California Verbal Learning Test. ART = Autobiographical Recollection Test. ASQ = Attentional Style Questionnaire. BIS-11 = Barratt Impulsiveness Scale. HADS = Hospital Anxiety and Depression Scale. GDS = Geriatric Depression Scale. Values are presented as *M* (SD) [min, max]. The column “n (YA/OA)” indicates the number of participants in each group (Younger Adults / Older Adults). For neuropsychological variables, extreme values (> 3 SD from the group mean, computed separately within each group and variable) were replaced with NA prior to statistical analyses. Group differences were assessed using independent-samples Welch *t*-tests (two-tailed) or with non-parametric Wilcoxon rank-sum test where normality assumptions were violated. Gender is reported as n (%) and compared using $\chi^2$ tests. Effect sizes are reported as Cohen’s *d* for parametric variables, as rank-biserial *r* when normality assumptions were violated, and as $\phi$ for gender. Significance levels are indicated as follows: $p < .05^*$ , $p < .01^{**}$ , $p < .001^{***}$ .

Episodic memory was assessed using the California Verbal Learning Test (CVLT; Delis et al., 2016). Autobiographical memory was measured using the Autobiographical Recollection Test (ART; Berntsen et al., 2019), which captures individual differences in richness and vividness of recollective experience. Executive functioning was evaluated through inhibitory control using the Victoria Stroop Test (Stroop, 1935; Spreen & Strauss, 1998) and working memory using both an N-Back task (Kirchner, 1958) and digit span measures from the Wechsler Adult Intelligence Scale (WAIS-III; Wechsler, 1997). Processing speed was assessed with the Digit Symbol Substitution Test (WAIS-III; Wechsler, 1997). Attentional orientation toward internal versus external focus was measured using the Attentional Style Questionnaire (ASQ; Van Calster et al., 2018). Trait impulsivity was assessed with the Barratt Impulsiveness Scale (BIS-11; Barratt, 1994; Baylé et al., 2000). Depression and anxiety were measured using the Hospital Anxiety and Depression Scale (HADS; Zigmond & Snaith, 1983) and the Geriatric Depression Scale (GDS; Yesavage et al., 1982).

Nominal associations between neuropsychological measures and implicit learning performance were observed in both groups, but none survived FDR correction. These associations are reported in full in Supplementary Table S2 and Supplementary Figure S1.

### 2.5. EEG recording and preprocessing

#### EEG acquisition

EEG signal was recorded continuously in a faraday cage (Tucker, 1993; GES 400 amplifier, EGI Hydrocel Geodesic Sensor Net, 128 Ag/AgCl dense array sensors), referenced (vertex reference Cz, ground reference CPz), and sampled at 1 kHz frequency with a 24-bit A/D. Impedances were kept under 100 kΩ, with a majority of electrodes below 50 kΩ (Ferree et al., 2001). Horizontal and vertical electrooculograms (EOG) were recorded during the session to improve artifact cleaning.

#### EEG preprocessing

EEG raw data were processed offline using Brainstorm (version 3.260407; Tadel et al., 2011) running under Matlab environment (Version: 24.1.0.2689473 (R2024a) Update 6; The MathWorks Inc., 2024). Data were band-pass filtered between 0.1 and 60 Hz and notch filtered at 50 Hz. Ocular and cardiac artifacts were detected and corrected using signal-space projection (SSP). The number of removed SSP components was comparable across recordings, with a mean of 4.40 (range: 1-9). Data were visually inspected, and noisy segments were rejected. Subjects were retained only if at least 75% of the recording remained artifact-free after rejection. EEG data were then re-referenced using the algebraic average of the left and right mastoid electrodes (Luck, 2014). Bad channels were identified and interpolated using spherical spline interpolation. On average, 5.71 ± 4.02 channels (4.46% of the 128 scalp electrodes; range: 0-20) were interpolated per participant based on visual inspection (no significant group difference; younger adults: 5.23 ± 3.33, 4.09%; older adults: 6.14 ± 4.55, 4.80%; Wilcoxon test, *p* =.566).

#### Epoching

EEG data were segmented time-locked to stimulus onset (i.e., onset of the visual stimulus). Epochs were extracted from-500 ms to 1000 ms relative to stimulus onset. Baseline correction was applied using the pre-stimulus interval (-500 to 0 ms). Only trials with correct responses were retained for subsequent analyses. After exclusion of incorrect trials, the total number of artifact-free epochs yielded on average 87.79 epochs (range: 68-95) for the familiar condition and 84.20 epochs (range: 65-94) for the novel condition per participant. Of note, these differences were statistically significant between conditions (paired *t*(65) = 5.28, *p* <.001) and between groups, with younger adults retaining more epochs than older adults for both familiar (younger adults: M = 89.00, SD = 3.67; older adults: M = 86.71, SD = 4.97; *t*(64) = 2.10, *p* =.040) and novel items (younger adults: M = 87.06, SD = 4.65; older adults: M = 81.66, SD = 6.17; *t*(64) = 3.98, *p* <.001).

Given the tendency of participants to give their answer before the response screen appeared, particularly in older adults, following analyses were then restricted to the 0-600 ms post-stimulus window to minimize potential contamination from motor-related activity.

While behavioral analyses were conducted on all four blocks separately to capture the full learning trajectory, this block-wise structure did not yield a sufficient number of artifact-free epochs per condition for robust time-frequency estimation. EEG analyses were therefore conducted on the first and last thirds of trials pooled across the four blocks, rather than block by block, allowing direct comparison of oscillatory activity between early and late learning stages while maintaining enough trials per condition. This selection resulted in 22 to 31 trials per condition and per participant within each learning stage. Stimulus positions (first vs. second presentation within a trial) were then merged by averaging across positions to isolate neural activity related to implicit encoding processing independently of stimulus order.

#### Time-frequency analysis

Time-frequency representations were computed using Morlet wavelet convolution (Tallon-Baudry et al., 1997; Cohen, 2014). Power estimates were obtained across predefined frequency bands (delta: 2-4 Hz; theta: 5-7 Hz; alpha: 8-12 Hz; beta: 13-29 Hz; gamma: 30-59 Hz). Spectral normalization was applied to reduce broadband spectral bias related to the 1/f structure of the EEG signal. Event-related synchronization/desynchronization (Clochon et al., 1996; Pfurtscheller, 1992) was calculated relative to the baseline period (-500 to 0 ms). Time-frequency maps were averaged per condition and per third of block before group-level analyses.

#### Cluster-based permutation tests

Condition differences (Familiar vs. Novel) in time-frequency power were assessed separately for each group (Young, Older) and for each third of block using two-tailed cluster-based permutation tests (1000 permutations; Maris & Oostenveld, 2007). Clusters were defined based on spatial adjacency (minimum three neighboring electrodes) and a cluster-forming threshold of *p* <.05. Statistical significance was assessed independently within each group.

To quantify task-related theta modulation for EEG-behavior analyses, a theta (5-7 Hz) Familiar-Novel contrast index was extracted for each participant by averaging power differences within the time-frequency window and electrode cluster identified in younger adults, separately for the early and late learning stages (corresponding to the first and last third of trials pooled across the four blocks). This procedure yielded two indices which were then used in correlation analyses with behavioral performance measures. Importantly, this cluster definition was held fixed for index extraction and was not used to re-test condition effects in the older group.

### 2.6. Statistical analyses

Statistical analyses were conducted in R (version 4.5.2; R Core Team, 2025-09-02, ucrt build, https://cran.r-project.org). Missing values were excluded at the variable level, while participants were retained unless all observations for a given analysis were missing. Observations exceeding ±3 standard deviations from the group mean were treated as outliers and excluded prior to inferential testing. Assumptions of normality and homogeneity of variance were assessed using Shapiro-Wilk and Levene’s tests, respectively before statistical analyses were applied.

Unless indicated otherwise, all results presented below are means ± standard deviation (SD). Due to significant differences between group, years of education were included in all between-group analyses as a covariate. Its influence is reported only when significant. To control for multiple comparisons, *p*-values were adjusted using the Benjamini-Hochberg false discovery rate (FDR) procedure within each family of tests. As an exception, *p*-values for correlations between theta dynamics and implicit learning performance were not adjusted for multiple comparisons due to the limited number of a priori individual hypotheses (as discussed by Rubin, 2024 and García-Pérez, 2023). Results were considered statistically significant at the *p* ≤ 0.05 level.

Two mixed-design ANCOVAs were conducted with Group (Young, Older) as the between-subject factor and years of education as a covariate: the first examined accuracy and response times during the implicit learning phase, with Condition (Familiar, Novel) and Block as within-subject factors; the second examined recognition accuracy, with Task (DIILT, Explicit Encoding) and Session (Immediate, Delayed) as within-subject factors. For the latter, a retention index (Δ = immediate - delayed) was additionally computed for each task and analyzed using the same ANCOVA structure.

To further characterize learning dynamics, additional indices were computed for each condition (Familiar, Novel), including Area Under the Curve (AUC), a differential learning index (delta Familiar-Novel) and Coefficient of Variation (CV). Group differences on these indices, as well as on task-related theta modulation, were assessed using Welch *t*-tests or Wilcoxon rank-sum tests. Associations between theta dynamics and behavioral performance were examined using partial correlations, controlling for years of education.

## 3. Results

### Younger adults more rapidly took advantage of the repetition of the novel implicit associations compared to older adults

The performance obtained during DIILT is illustrated in Figure 2A. Accuracy analysis revealed significant main effects of Group (*F*(1, 59) = 14.77, *p* <.001, ηp^2^ =.200) and Condition (*F*(1, 59) = 65.42, *p* <.001, ηp^2^ =.526), as well as a significant Group x Condition interaction (*F*(1, 59) = 13.52, *p* <.001, ηp^2^ =.186). Critically, post hoc comparisons revealed that older adults showed consistently higher accuracy for familiar compared to novel items across all four blocks (all *p*s <.001), whereas in younger adults, this advantage was only significant at Block 1 (*p* =.018), with performance on familiar and novel items converging from Block 2 onwards (all *p*s >.123). Although the Group x Block interaction did not reach significance (*F*(2.93, 172.94) = 1.26, *p* =.290), planned contrasts from the model confirmed a significant accuracy improvement for novel items between Block 1 and Block 4 in younger adults (*t*(59) =-2.64, *p* =.011), but not in older adults (*t*(59) =-0.93, *p* =.356), indicating that overall learning gains across the task were specific to the younger group.

**Figure 2.**
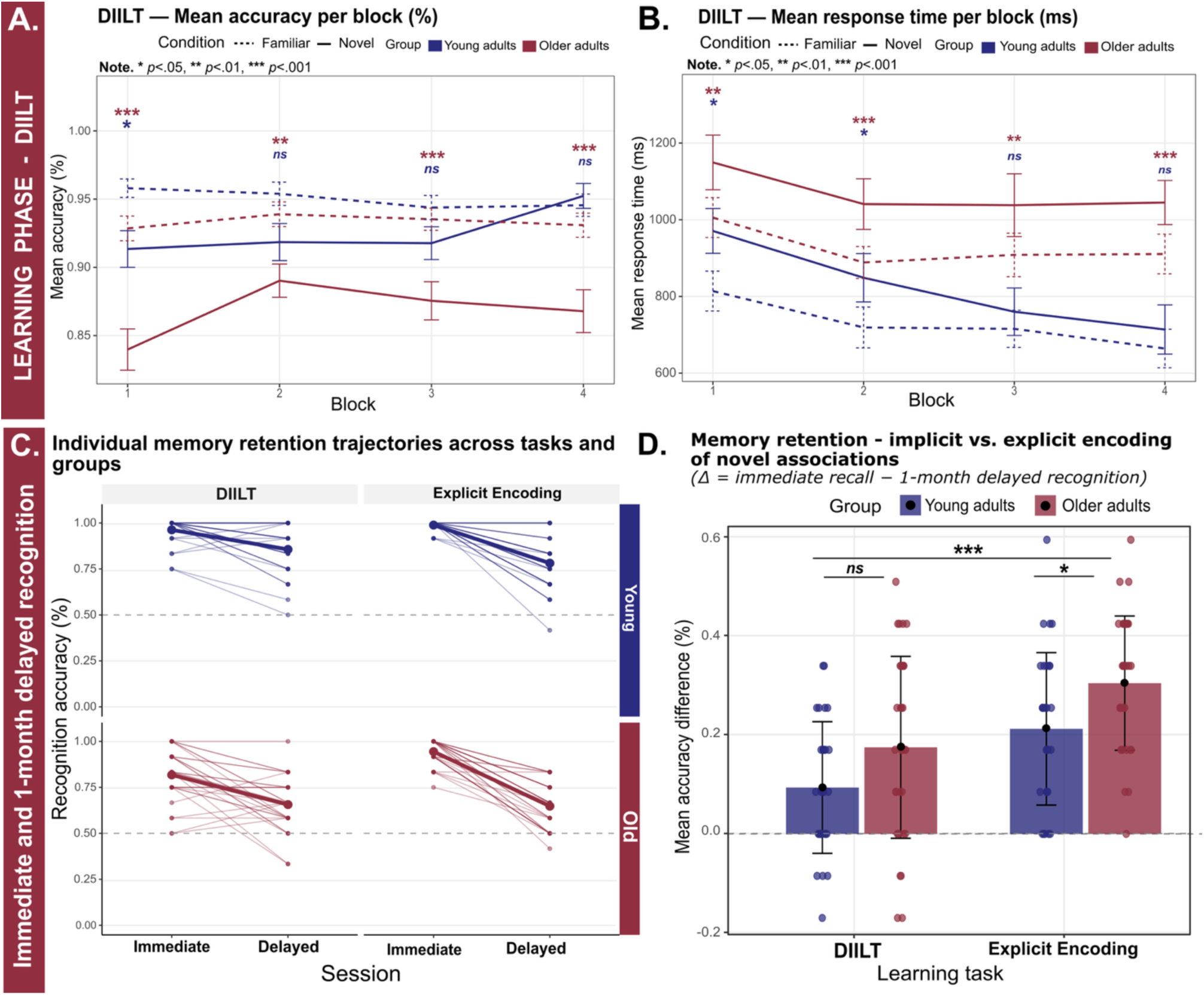
DIILT learning phase performance and memory retention across groups. (A) and (B): Mean accuracy (%) and mean response time (ms) during the implicit learning phase across blocks and conditions for younger (dark blue) and older adults (burgundy). Dashed lines represent familiar items, and solid lines represent novel items. Error bars indicate standard error. Significance indicators reflect post hoc comparisons (familiar vs. novel) within each group at each block: * *p* <.05, ** *p* <.01, *** *p* <.001. (C) Memory retention over one month for implicit and explicit learning tasks. Bars represent mean accuracy difference (Δ = immediate - delayed recognition) for younger (dark blue) and older adults (burgundy). Error bars indicate standard deviation. Positive values indicate forgetting, with a delta score closer to zero meaning greater retention over time. (D) Individual memory retention trajectories from immediate to delayed recognition for implicit (DIILT) and explicit encoding tasks, separately for younger (dark blue) and older adults (burgundy). Each line represents one participant. Thicker lines represent group means. The dashed horizontal line indicates chance level (50%). The figure illustrates the variability in individual forgetting patterns across groups and tasks.

The same pattern was observed for response times, with a significant main effect of Group (*F*(1, 60) = 13.20, *p* <.001, ηp^2^ =.180), Condition (*F*(1, 60) = 37.56, *p* <.001, ηp^2^ =.385), and Block (*F*(2.10, 126.18) = 13.51, *p* <.001, ηp^2^ =.184), indicating faster responses for familiar items and a general speed improvement across blocks for both groups. The Group x Condition interaction did not reach significance (*F*(1, 60) = 3.31, *p* =.074, ηp^2^ =.052). Post-hoc comparisons revealed that younger adults showed slower responses for novel compared to familiar items at Block 1 (*p* =.016) and Block 2 (*p* =.043) only, whereas older adults showed this difference consistently across all blocks (all *p*s ≤.003; Figure 2B). Planned comparisons revealed significant speed improvements between Block 1 and Block 4 in younger adults for both novel (*t*(60) = 3.99, *p* <.001) and familiar items (*t*(60) = 2.84, *p* =.006). In older adults, no significant speed improvement was observed for either novel (*t*(60) = 1.66, *p* =.102) or familiar items (*t*(60) = 1.93, *p* =.058).

### Learning indices confirm faster novel association acquisition in younger adults

Younger adults showed significantly higher AUC for novel items compared to older adults (*W* = 791.5, *p* <.001, *p*FDR =.002, *r* =.51), reflecting greater overall accuracy for novel associations across the learning phase. Moreover, the Familiar-Novel accuracy differential at Block 4 was significantly larger in older adults (*t*(57.75) =-3.51, *p* <.001, *p*FDR =.002, *d* =-0.85), indicating a more pronounced reliance on familiar items relative to novel ones that persisted until the end of the learning phase. A trend-level difference was also observed for the Familiar-Novel accuracy differential at Block 1 (*t*(60) =-2.14, *p* =.036, *p*FDR =.072), suggesting this pattern was already present at the beginning of the learning phase. However, no significant group differences were observed for the Coefficient of Variation (novel: *p* =.060; familiar: *p* =.751, all *p*FDR >.13), indicating similar intra-individual consistency in learning trajectories across groups.

### Both groups successfully retained novel associations over one month, with implicit learning showing better long-term memory traces than explicit encoding

The recognition accuracy analysis revealed significant main effects of Group (*F*(1, 59) = 27.72, *p* <.001, ηp^2^ =.320) and Session (*F*(1, 59) = 203.89, *p* <.001, ηp^2^ =.776), with younger adults showing consistently higher recognition accuracy and both groups showing a significant decline from immediate to delayed recognition. The main effect of Task was not significant (*F*(1, 59) = 1.45, *p* =.234). Significant interactions were found for Group x Session (*F*(1, 59) = 7.07, *p* =.010, ηp^2^ =.107) and Task x Session (*F*(1, 59) = 20.06, *p* <.001, ηp^2^ =.254). The Group x Task interaction (*F*(1, 59) = 3.90, *p* =.053, ηp^2^ =.062) and the three-way interaction (*p* =.765) did not reach significance. Post hoc comparisons confirmed that both groups showed a significant reduction in recognition accuracy from immediate to delayed session across all conditions (all *ps* ≤.008), reflecting forgetting over one month. Nevertheless, performance remained above chance at both immediate and delayed recognition across both tasks (all *t*s > 6.31, all *p*s <.001, all *p*FDR <.001). Importantly, younger adults showed significantly better performance than older adults at both immediate and delayed recognition, across both implicit (DIILT: immediate, M_young =_.96, SD =.08 vs. M_old =_.82, SD =.14, *p* =.002; delayed, M_young =_.86, SD =.14 vs. M_old =_.65, SD =.14, *p* <.001) and explicit tasks (EE: immediate, M_young =_.99, SD =.03 vs. M_old =_.94, SD =.08, *p* =.005; delayed, M_young =_.78, SD =.15 vs. M_old =_.64, SD =.13, *p* <.001). Additional comparisons between tasks revealed that this advantage for DIILT differed across groups and sessions: in younger adults, immediate recognition did not differ between tasks (*p* =.115), but most importantly, implicit recognition was significantly higher than explicit one at the delayed session (*p* =.039). Notably, in older adults, associations learned during the explicit task yielded significantly higher immediate recognition than during the implicit task (*p* <.001), whereas no difference between tasks was observed at the delayed session (*p* =.459) (Figure 2C).

The mixed ANCOVA on delta scores (Figure 2D) revealed a significant main effect of Task (*F*(1, 59) = 20.06, *p* <.001, ηp^2^ =.254), confirming greater forgetting for explicitly encoded than implicitly learned associations in both groups. A significant main effect of Group was also observed (*F*(1, 59) = 7.07, *p* =.010, ηp^2^ =.107), with older adults showing overall greater forgetting than younger adults. The Group x Task interaction was not significant (*p* =.765).

### Task-related theta modulation during implicit learning (DIILT) differs between younger and older adults

To compare oscillatory activity between early and late learning stages, EEG analyses focused on the first and last third of trials pooled across the four learning blocks, ensuring a sufficient number of artifact-free trials per condition for time-frequency estimation (see Methods). In younger adults (Figure 3A), a significant negative cluster was identified at the early stage of learning (*p* =.002), reflecting greater theta power (5-7 Hz) for novel compared to familiar items over posterior electrodes (161-499 ms; electrodes PO3, P1, PZ, PO7, PPO1h, POO1, POZ, OZ, POO2, PPO2h, P2, O2, PO8, PO4). This cluster dissipated in the late stage of learning, reflecting the absence of difference in terms of brain activity between novel items compared to familiar ones. Conversely, in older adults (Figure 3B), no cluster was found at the early stage of learning. In the late learning stage, a significant positive broadband cluster (alpha-beta-gamma, 8-30+ Hz) was found (*p* =.002), reflecting overall greater power for familiar compared to novel items over a widespread scalp distribution (476-600 ms; n = 79 electrodes).

**Figure 3.**
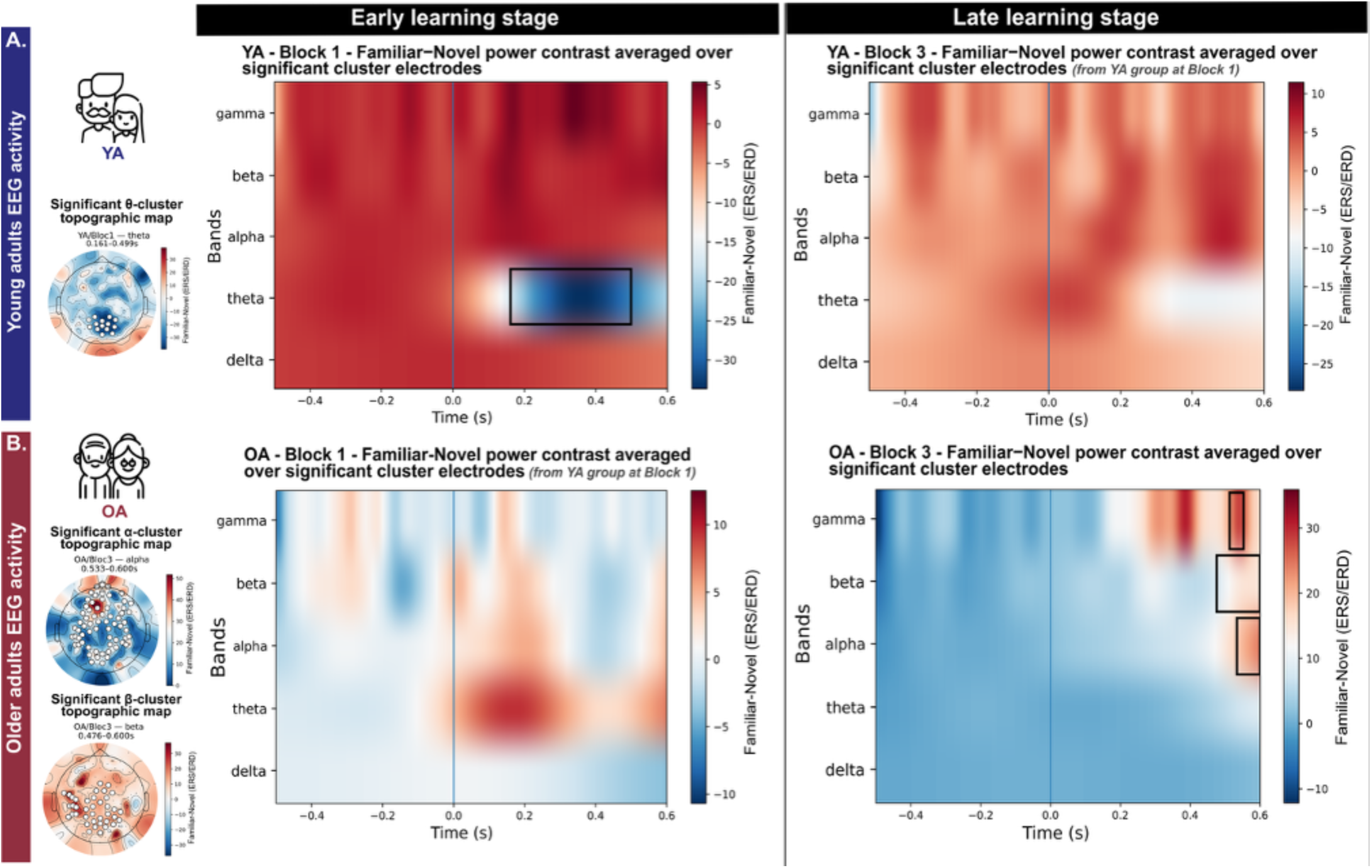
Task-related oscillatory activity (Familiar-Novel) during the DIILT learning phase. Time-frequency representations of the Familiar-Novel power contrast averaged over cluster electrodes, for younger (A; top) and older adults (B; bottom) at the early (left) and late (right) stages of learning. Black rectangles indicate significant clusters identified by cluster-based permutation tests (*p* <.05, 1,000 permutations). Topographic maps display the spatial distribution of the significant cluster for each condition. For visualization purposes, when no cluster was found, condition differences (Familiar-Novel) were averaged within the significant time-frequency window and sensor cluster identified in the younger adults’ group at Block 1, used as a reference for the time-frequency signature of theta modulation, in order to visualize how this modulation departs from this reference across groups and learning stages, using MNE-Python (Gramfort, 2013). (A) *Left panel*, younger adults, early learning stage: significant negative theta cluster (Familiar < Novel, *p* =.002, 161–499 ms) over posterior electrodes, reflecting greater theta synchronization for novel items. *Right panel*, younger adults, late learning stage: no significant cluster. (B) *Left panel*, older adults, early learning stage: no significant cluster. *Right panel*, older adults, late learning stage: significant positive broadband cluster spanning alpha, beta, and gamma frequencies (Familiar > Novel, *p* =.002, 476–600 ms) over a widespread scalp distribution. Color scale represents Familiar-Novel power difference (ERS/ERD).

### Task-related theta modulation during implicit learning differs between groups and relates to subsequent memory performance

During the early learning stage, older adults showed significantly higher theta contrast values compared to younger adults (*t*(62.72) =-3.55, *p* <.001, *p*FDR =.004, *d* =-.86), indicating greater theta power for familiar relative to novel items at the beginning of the task in older adults (Figure 4A). This difference in theta contrast then disappeared for the late learning stage between groups (*p* =.347).

**Figure 4.**
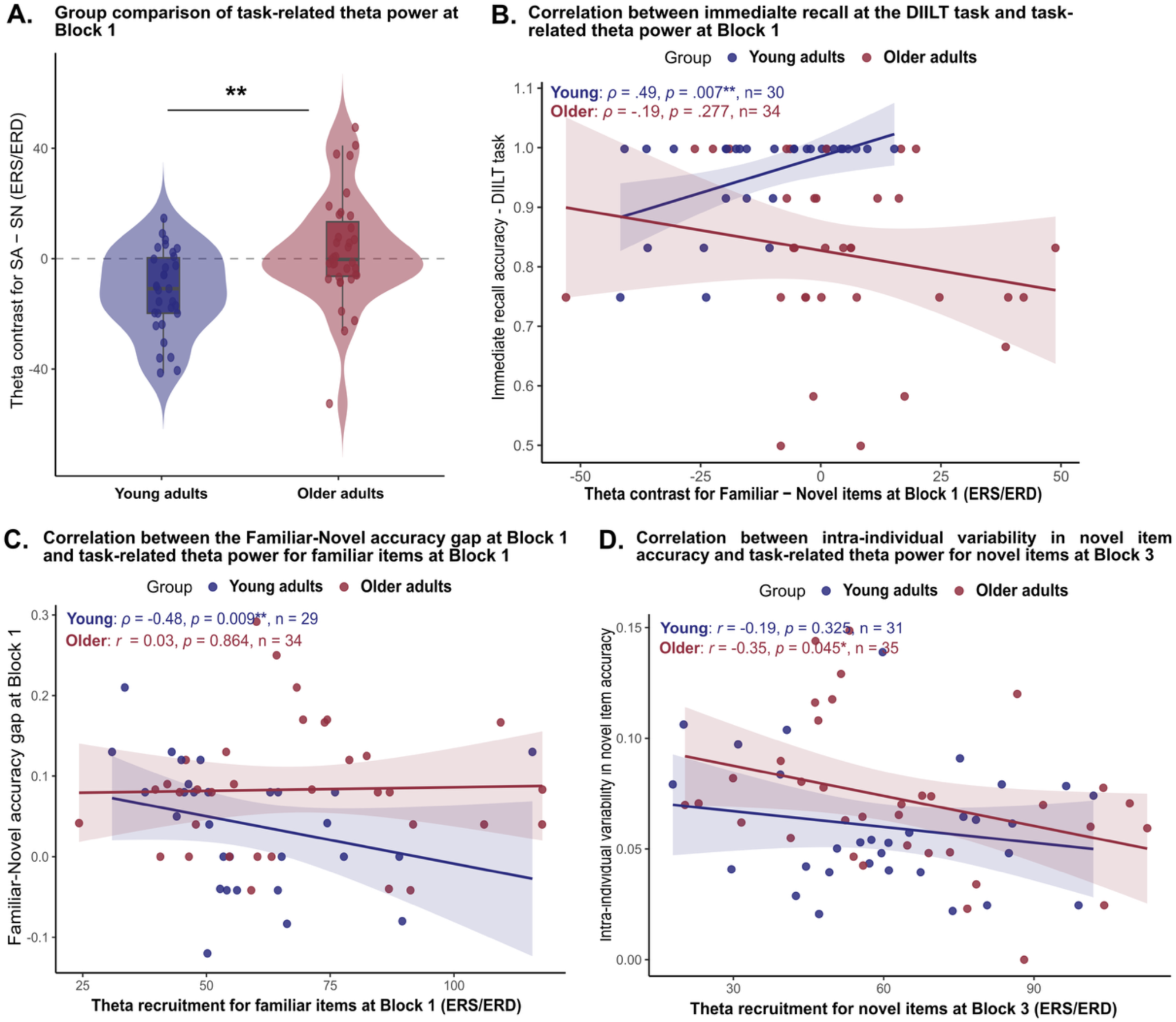
**Theta modulation for novel items during implicit learning predicts memory in younger but not older adults**. (A) Violin plots showing the distribution of the theta power contrast (Familiar-Novel) at the early learning stage for younger (dark blue) and older adults (burgundy). Negative values indicate greater theta power for novel than familiar items. The dashed line indicates zero. Older adults showed significantly higher theta contrast values compared to younger adults (*p* <.001, *p*FDR =.004). (B) Scatterplot with linear fit showing the relationship between theta contrast (Familiar - Novel, dB) at Block 1 (early learning stage) and immediate recognition accuracy (proportion correct) for novel items, controlling for years of education. A significant positive association was observed in younger adults (*ρ* =.49, *p* =.007), but not in older adults (*ρ* =-.19, *p* =.277). (C) Scatterplot with linear fit showing the relationship between theta power for familiar items at Block 1 (early learning stage) and the Familiar-Novel accuracy gap (proportion correct) at Block 1, controlling for years of education. A negative association was observed in younger adults (*ρ* =-.48, *p* =.009), but not in older adults (*r* =.03, *p* =.864). (D) Scatterplot with linear fit showing the relationship between theta power for novel items at Block 3 (late learning stage) and intra-individual variability in novel item accuracy (coefficient of variation) in older adults (*r* =-.35, *p* =.045) and younger adults (*r* =-.19, *p* =.325), controlling for years of education. **Note**. For panels B, C, D, *r, ρ* and *p* values correspond to Spearman or Pearson correlations depending on normality. Given the small number of a priori hypothesized theta-behavior associations, *p*-values for theta x implicit learning performance correlations were not FDR-corrected. Dark blue = younger adults; burgundy = older adults. Significance indicators are indicated as follows: *p* <.05*, *p* <.01**, *p* <.001***.

In younger adults, higher theta contrast at Block 1 (early learning stage) was positively correlated with immediate recognition accuracy for novel items learned during the implicit task (*ρ* =.49, *p* =.007; Figure 4B). In older adults, no significant association between theta contrast and DIILT performance was observed. In younger adults, higher theta for familiar items at Block 1 was negatively associated with the Familiar-Novel accuracy gap at Block 1 (*ρ* =-.48, *p* =.009; Figure 4C), while higher theta for novel items at Block 3 (late learning stage) was negatively associated with immediate DIILT recognition (*ρ* =-.40, *p* =.031). Critically, in older adults, higher theta for novel items at Block 3 was negatively associated with intra-individual variability in novel item accuracy (*r* =-.35, *p* =.045; Figure 4D). Additionally, theta for novel items at Block 1 was positively associated with processing speed in this group (*r* =.36, *p* =.036).

## 4. Discussion

The present study aimed to characterize implicit learning across age groups using an EEG-adapted Fast Mapping paradigm (the DIILT), focusing on behavioral performance and task-related theta dynamics over the course of learning. At the behavioral level, we expected that both groups would successfully encode novel associations implicitly, but that older adults would show less improvement across repeated exposures, resulting in attenuated gains in novel item accuracy across blocks. We also hypothesized that younger adults would exhibit increased theta activity for novel associations that would decrease as learning stabilizes, whereas older adults were expected to show attenuated or redistributed theta modulation. Finally, we expected that theta dynamics during encoding would relate to subsequent memory performance, with the nature of this association differing between age groups, reflecting a reorganization of the neural substrates supporting implicit learning with age. Overall, our results partially validate our hypotheses. We confirmed age-related differences in novel association acquisition, with younger adults showing faster and more stable integration of novel items, while both groups successfully formed durable memory traces. Task-related theta modulation differed between groups and related differentially to memory performance.

At the behavioral level, younger adults rapidly took advantage of repeated exposures to referent-image associations when performing the DIILT task. The progressive reduction in reaction times across repetitions, together with the observation that reaction times in the novel condition gradually approached those observed in the familiar condition, suggests the acquisition of associative links between the referents and their corresponding images. Our results are in line with current literature on Fast Mapping and statistical learning in healthy adults, showing an effective integration of novel implicit associations into existing knowledge networks when the task promotes independence and differentiation (Merhav et al., 2015; Sharon et al., 2011). By contrast, older adults maintained a persistent reliance on familiar associations throughout the task and showed less improvement than younger adults for novel items beyond the early learning phase. This pattern is consistent with a broader literature suggesting that the processing of novelty, a critical trigger for memory encoding (Tulving & Kroll, 1995), can be altered with age, more markedly so in pathological aging (Bastin et al., 2019), although this decline is not systematic and can be relatively preserved in cognitively high-performing older adults (Daffner et al., 2006). Nonetheless, it is worth noting that older adults’ overall accuracy for novel items remained high from Block 2 onwards, suggesting that the observed group differences reflect a slower learning trajectory rather than an inability to perform the task. Indeed, successful task performance does not necessarily imply the retention of novel associations across trials, but rather correct inference from the disjunctive structure on each trial independently. These results are consistent with Fast Mapping studies comparing younger and older adults, which have generally shown that older adults fail to fully close the performance gap with novel items despite equivalent task exposure (Greve et al., 2014), and with broader findings on implicit learning in aging suggesting that while novel association formation remains available, its efficiency and extent are reduced with age (Schevenels et al., 2023; Wagnon et al., 2019). They also further extend findings from statistical learning research, where age-related differences in overall performance have been reported alongside slower behavioral adaptation in older adults (Bulgarelli et al., 2021; Herff et al., 2020).

Several factors may account for this behavioral dissociation. Younger adults succeeded in inferring novel labels from their co-occurrence with known referents and efficiently allocated their attention toward novelty, whereas older adults appeared to rely more heavily on familiar associations. This persistent reliance on familiar items in older adults, alongside limited improvement for novel associations, may reflect reduced availability of the attentional and inferential resources required to engage the exclusion-based mechanism the task was designed to elicit. Consistent with Zaiser et al. (2022), who showed that semantic feature overlap between known and novel items modulates the extent of rapid cortical integration, older adults may have relied more heavily on pre-existing semantic knowledge due to reduced capacity to engage novelty-driven processing. Additionally, the duration of the DIILT task, which corresponds to approximately 40 minutes of continuous cognitive engagement, may have also contributed to differential performance across groups. Indeed, older adults are known to be more susceptible to cognitive fatigue during sustained cognitive tasks, which can impair attentional resources and accuracy over time (Bouche et al., 2025; Gilsoul et al., 2024). This is consistent with the pattern observed here, where descriptively older adults appeared to show an initial improvement between blocks 1 and 2 but failed to maintain or build upon this gain in later blocks; a trajectory reminiscent of what Schevenels et al. (2021) described as performance under high attentional or strategic demands.

With regard to memory retention over one month, both groups successfully encoded novel associations through both learning procedures, confirming that implicit encoding produces durable memory traces even in older adults. As expected, younger adults showed better recognition than older adults at both immediate and delayed sessions across both tasks. Rather than reflecting memory decline *per se*, this pattern may reflect differences in the quality of semantic integration during the learning phase. Younger adults successfully integrated novel associations into existing semantic networks, as suggested by the convergence of novel and familiar item processing from Block 2 onwards, whereas the persistent reliance on familiar items in older adults suggests less complete neocortical integration, potentially resulting in less durable memory traces (Coutanche & Thompson-Schill, 2014; Merhav et al., 2015; Sharon et al., 2011).

Moreover, the pattern of retention consistently favored implicitly learned associations in both groups. In older adults, despite lower immediate performance for implicitly than explicitly learned associations, implicitly learned associations were better preserved over one month. In younger adults, immediate performance was equivalent across tasks, but implicitly learned associations again showed superior long-term retention. This dissociation between immediate and long-term performance echoes previous Fast Mapping studies showing greater long-term stability for associations learned through Fast Mapping compared to explicitly encoded associations (Chen et al., 2023; Li et al., 2020), and is consistent with a broader literature showing that while explicit memory declines with age, implicit memory measures, including implicit associative memory, tend to remain comparatively stable (Davis et al., 2021; Fleischman et al., 2004). Together, this convergence across age groups points toward a more general encoding route for implicitly learned associations, distinct from the mechanisms supporting explicit memory, which rely more heavily on the hippocampus and become increasingly vulnerable over the lifespan (for a review on hippocampus and aging, see Bettio et al., 2017). One possibility, still debated in the Fast Mapping literature (Cooper et al., 2019b), is that this route partly relies on rapid neocortical plasticity less dependent on the hippocampus (Atir-Sharon et al., 2015), a more general mechanism that could be comparatively preserved with age.

At the neural level, task-related oscillatory dynamics during implicit learning differed between age groups. To our knowledge, this study provides the first direct comparison of oscillatory dynamics during adapted Fast Mapping encoding between younger and older adults, thereby extending previous EEG evidence limited to neural responses measured before and after learning (Shtyrov, 2011; Shtyrov et al., 2021, 2022; Vasilyeva et al., 2019). In younger adults, greater theta power for novel items compared to familiar items was observed over parietal electrodes during the early learning phase, consistent with the role of theta oscillations in associative encoding and successful memory formation (Backus et al., 2016; Herweg et al., 2020; Lega et al., 2012). By the end of the learning blocks, this theta activity decreased, mirroring the behavioral convergence between novel and familiar item accuracy observed from Block 2 onwards, and indicating that theta engagement for novel items could reflect an early binding process that stabilizes as associations become stronger.

In older adults, no such theta modulation was observed for novel items at the beginning of learning. Importantly, a substantial number of older adults seemed to exhibit theta recruitment levels comparable to those of younger adults. Notably, theta recruitment for novel items at the end of learning was negatively associated with intra-individual variability in novel item accuracy in older adults, suggesting that those who maintained stronger theta engagement showed more consistent encoding across the task. This observation is consistent with the age-related reorganization of the oscillatory mechanisms supporting semantic learning (Chiang et al., 2023) and, importantly, with the concept of neural maintenance in aging, whereby some individuals preserve oscillatory dynamics similar to younger adults, independently of compensatory recruitment, as described in recent M/EEG frameworks of cognitive aging (Jauny et al., 2022).

More broadly, theta dynamics during encoding related differentially to subsequent memory performance across age groups. In younger adults, greater theta engagement for novel items at the beginning of learning positively predicted immediate recognition accuracy for implicitly learned associations, suggesting that early theta engagement reflects effective binding of novel information through hippocampo-cortical feedback loops (Backus et al., 2016; Klimesch, 1999). Although scalp EEG primarily captures neocortical theta (∼6 Hz), recent intracranial evidence suggests that human hippocampal encoding is organized by slower oscillations (∼2 Hz) that are coupled with neocortical theta during learning, and that stronger hippocampal slow-oscillatory activity predicts better memory formation (Causse et al., 2026). The posterior theta we observe may thus reflect the neocortical signature of this hippocampo-cortical coupling. In older adults, beyond the maintenance-related association described above, theta engagement during implicit learning was positively associated with processing speed rather than memory performance, suggesting that theta engagement in this group may reflect individual levels of available cognitive resources rather than memory-specific binding. This result is consistent with previous work showing that implicit encoding performance is more strongly coupled with attentional and executive resources in older adults compared to younger adults (Chiang et al., 2023; Kontaxopoulou et al., 2017) and suggests that the efficiency of implicit encoding in older adults may vary as a function of individual cognitive resources. Together, these dissociations suggest an age-related shift in the functional role of theta during implicit encoding: from a memory-specific binding mechanism in younger adults to a possible marker of individual cognitive maintenance in older adults.

This interpretation is further supported by the neuropsychological correlates of learning performance, which also differed between age groups (see Supplementary Table S1 and Figure S1 for full results). In younger adults, better working memory capacity was associated with higher accuracy for novel associations and a smaller reliance on familiar items at the end of learning, providing behavioral support for the role of working memory in Fast Mapping performance (Greve et al., 2014). In older adults, better processing speed was associated with higher immediate recognition for implicitly learned associations, converging with the oscillatory finding that theta engagement for novel items relates to processing speed in this group. Additionally, older adults with better episodic memory showed less reliance on familiar items from the outset of learning, consistent with the idea that intact episodic memory resources reduce over-reliance on pre-existing semantic knowledge during implicit encoding (Umanath & Marsh, 2014). Together, these age-specific associations suggest that implicit encoding relies on different cognitive substrates across the lifespan, and that implicit encoding may be particularly sensitive to individual differences in cognitive preservation in older adults.

This study provides one of the first direct comparisons between younger and older healthy adults during the implicit encoding of new information and its behavioral and neural age-related differences, but there are some limitations to acknowledge. First, the two groups differed significantly in years of education, which was included as a covariate in all between-group analyses but could not be fully controlled at the recruitment stage. However, its influence was non-significant across our results. Second, the relatively modest sample size limits the examination of potential subgroups within the older adult population, such as individuals showing maintenance versus decline patterns in oscillatory dynamics. Future work with larger samples would allow for a more fine-grained characterization of inter-individual variability in implicit encoding across aging. Third, the behavioral pattern observed here, with older adults showing persistent reliance on familiar associations while younger adults efficiently engaged novel item processing, could be interpreted in terms of differential strategy use at encoding, in line with age-related differences observed in strategy repertoire, distribution, execution, and selection (Lemaire, 2010). However, as the present design does not include a direct measure of encoding strategy, this interpretation remains speculative, and disentangling genuine age-related differences in implicit encoding from differences in task strategy remains methodologically challenging. As Cooper et al. (2019) have argued, inconsistencies across Fast Mapping studies may partly reflect variability in what the task actually measures, rather than true differences in implicit learning capacity. Future studies incorporating think-aloud protocols or strategy questionnaires and manipulating task demands and encoding conditions across age groups, will be needed to clarify this interpretation. Additionally, the present study does not include measures of cognitive fatigue, which is proposed here to have contributed to the performance decline observed in older adults during later learning blocks. Future work adding subjective fatigue ratings or physiological measures could more directly test this hypothesis. Finally, our DIILT task constitutes an approximation of classical Fast Mapping paradigm adapted to meet the constraints of EEG recording. While this adaptation was necessary to capture oscillatory dynamics unfolding during learning, interpretations of the present findings in relation to the broader Fast Mapping literature should take this methodological difference into account.

In conclusion, these findings have implications for current frameworks of implicit learning and cognitive aging (e.g., Park & Reuter-Lorenz, 2009; Hedden & Gabrieli, 2004; Schevenels et al., 2021), as they contribute to characterizing the behavioral and neural mechanisms through which the aging brain forms new memory traces without deliberate intent. By recording oscillatory dynamics during Fast Mapping encoding itself, rather than before or after learning, and assessing memory retention over a one-month delay, this study addresses two critical gaps in the literature and provides a more ecologically valid window into how implicit learning unfolds and persists in everyday life. Our results demonstrate that implicit encoding remains available in older adults but relies on reorganized oscillatory dynamics, extending our understanding of how theta-dependent encoding mechanisms are reshaped across the lifespan. Furthermore, we show that implicit encoding not only produces more stable long-term memory traces than explicit encoding in both age groups, but also that this advantage is supported by distinct oscillatory mechanisms that are reorganized with age. Importantly, the association between theta dynamics and individual cognitive resources in older adults further suggests that implicit encoding may be sensitive to inter-individual variability in cognitive preservation in aging, with potential implications for the design of ecologically valid memory interventions in aging populations.

Future work combining multimodal neuroimaging will be needed to directly test whether the greater long-term stability associations learned through Fast Mapping reflects a more direct form of neocortical integration, and to better precise the structural and functional underpinnings of the oscillatory reorganization observed here.

## Ethical approval

The participants provided their written informed consent to participate in this study and the French ethics committee (Comité de Protection des Personnes, CPP; ID-RCB: 2024-A00167-40) approved all study procedures.

## Funding

This work was supported by the French National Research Agency (ANR) as part of the PRC ULIS project (ANR-22-CE28-0029).

## CRediT authorship contribution statement

### Florentine Fricker

Writing - Original Draft, Formal analysis, Visualization, Data Curation, Investigation, Methodology, Conceptualization, Writing - Review & Editing**. Vincent Dornier:** Conceptualization, Writing - Review & Editing**. Giulia Buzi:** Investigation, Writing - Review & Editing**. Maelys Morantin:** Investigation, Writing - Review & Editing**. Franck Doidy:** Investigation, Writing - Review & Editing**. Patrice Clochon:** Resources, Writing - Review & Editing**. Annabelle Goujon:** Conceptualization, Funding acquisition, Writing - Review & Editing**. Emmanuel J. Barbeau:** Supervision, Conceptualization, Funding acquisition, Writing - Review & Editing**. Thomas Hinault:** Writing - Original Draft, Supervision, Methodology, Conceptualization, Validation, Funding acquisition, Writing - Review & Editing

### Declaration of Competing Interest

None.

## Notes

### Competing Interest Statement

The authors have declared no competing interest.

