## Supplementary file for "How aging reshapes implicit learning: behavioral and oscillatory evidence from an EEG-adapted Fast Mapping paradigm"

### Statistical analyses

**Table S1. Complete DIILT item and question list across block.**

*Familiar associations.* Twelve items were selected and consisted in familiar fruits (banana, lemon, apple and strawberry), objects (mug, pan) and animals (duck, dog, cat, sheep, turtle, and butterfly). Four additional familiar items (crab, shrimp, chicken, starship) were included as potential referent but were never directly referenced in the questions.

*Novel associations.* Two lists of twelve novel items were created including novel objects (Loppe, Thiou, Poimon and Apoite), animals (Bardon, Edoime, Madrin, Plaime, Possecale, Rapare, Civane, Cratorre, Leipe, Phot, Setrin, and Tosse) and fruits (Crute, Musse, Nuraque, Toupe, Pesse, Guion, Caralle, and Brime), each associated with a pseudoword.

Familiar items were identical across both counterbalancing lists. Novel items were distributed across two lists (Even and Odd) counterbalanced across participants. Each item appeared twice per block with two different questions (one requiring a "yes" response and one requiring a "no" response). Block assignment reflects the sequential presentation order across the four blocks (two questions per item per block). Questions were generated using synthetic speech (TTSMaker; https://ttsmaker.com/; Dore's voice) and lasted approximately one second on average. A $t$-test on the number of letters per question revealed no difference in length between novel and familiar items ($t(262.99) = 0.82$, $p = .41$, n = 384).

| List | Condition | Item (English) | Block | Question (French) | Question (English) | Expected response |
|---|---|---|---|---|---|---|
| Referent | Familiar | Duck | 1 | Est-ce que ce canard a la tête verte ? | Does this duck have a green head? | yes |
| Referent | Familiar | Duck | 1 | Est-ce que ce canard a la tête marron ? | Does this duck have a brown head? | no |
| Referent | Familiar | Duck | 2 | Est-ce que ce canard a des plumes ? | Does this duck have feathers? | yes |
| Referent | Familiar | Duck | 2 | Est-ce que ce canard est en train de nager ? | Is this duck swimming? | no |
| Referent | Familiar | Duck | 3 | Est-ce que ce canard a des palmes ? | Does this duck have webbed feet? | yes |
| Referent | Familiar | Duck | 3 | Est-ce que ce bec du canard pointe vers la droite ? | Is the duck's beak pointing to the right? | no |
| Referent | Familiar | Duck | 4 | Est-ce que ce canard a le bec jaune ? | Does this duck have a yellow beak? | yes |
| Referent | Familiar | Duck | 4 | Est-ce que ce canard est en train de voler ? | Is this duck flying? | no |
| Referent | Familiar | Saucepan | 1 | Est-ce que cette casserole a un manche ? | Does this saucepan have a handle? | yes |
| Referent | Familiar | Saucepan | 1 | Est-ce que cette casserole a deux poignées ? | Does this saucepan have two handles? | no |
| Referent | Familiar | Saucepan | 2 | Est-ce que cette casserole est noire ? | Is this saucepan black? | yes |
| Referent | Familiar | Saucepan | 2 | Est-ce que cette casserole est pleine ? | Is this saucepan full? | no |
| Referent | Familiar | Saucepan | 3 | Est-ce que cette casserole est vide ? | Is this saucepan empty? | yes |
| Referent | Familiar | Saucepan | 3 | Est-ce que cette casserole est marron ? | Is this saucepan brown? | no |
| Referent | Familiar | Saucepan | 4 | Est-ce que cette casserole est ronde ? | Is this saucepan round? | yes |
| Referent | Familiar | Saucepan | 4 | Est-ce que cette casserole est carrée ? | Is this saucepan square? | no |

| | | | | | | |
|---|---|---|---|---|---|---|
| Referent | Familiar | Cat | 1 | Est-ce que ce chat vous regarde ? | Is this cat looking at you? | yes |
| Referent | Familiar | Cat | 1 | Est-ce que ce chat est tigré ? | Is this cat tabby-striped? | no |
| Referent | Familiar | Cat | 2 | Est-ce que ce chat a des pattes ? | Does this cat have paws? | yes |
| Referent | Familiar | Cat | 2 | Est-ce que ce chat est blanc ? | Is this cat white? | no |
| Referent | Familiar | Cat | 3 | Est-ce que ce chat est gris ? | Is this cat grey? | yes |
| Referent | Familiar | Cat | 3 | Est-ce que ce chat est en train de courir ? | Is this cat running? | no |
| Referent | Familiar | Cat | 4 | Est-ce que ce chat a une queue ? | Does this cat have a tail? | yes |
| Referent | Familiar | Cat | 4 | Est-ce que ce chat est roux ? | Is this cat ginger? | no |
| Referent | Familiar | Dog | 1 | Est-ce que ce chien est face à vous ? | Is this dog facing you? | yes |
| Referent | Familiar | Dog | 1 | Est-ce que ce chien a les poils ras ? | Does this dog have short fur? | no |
| Referent | Familiar | Dog | 2 | Est-ce que ce chien a des poils clairs ? | Does this dog have light-colored fur? | yes |
| Referent | Familiar | Dog | 2 | Est-ce que ce chien a des tâches ? | Does this dog have spots? | no |
| Referent | Familiar | Dog | 3 | Est-ce que ce chien a une grande queue ? | Does this dog have a long tail? | yes |
| Referent | Familiar | Dog | 3 | Est-ce que ce chien est en train de courir ? | Is this dog running? | no |
| Referent | Familiar | Dog | 4 | Est-ce que ce chien tire la langue ? | Is this dog sticking out its tongue? | yes |
| Referent | Familiar | Dog | 4 | Est-ce que ce chien est allongé ? | Is this dog lying down? | no |
| Referent | Familiar | Lemon | 1 | Est-ce que ce citron est jaune ? | Is this lemon yellow? | yes |
| Referent | Familiar | Lemon | 1 | Est-ce que ce citron est vert ? | Is this lemon green? | no |
| Referent | Familiar | Lemon | 2 | Est-ce que ce citron a des feuilles ? | Does this lemon have leaves? | yes |
| Referent | Familiar | Lemon | 2 | Est-ce que ce citron a des rayures ? | Does this lemon have stripes? | no |
| Referent | Familiar | Lemon | 3 | Est-ce qu'il y a plusieurs citrons ? | Are there multiple lemons? | yes |
| Referent | Familiar | Lemon | 3 | Est-ce que ce citron est accroché à son arbre ? | Is the lemon hanging on its tree? | no |
| Referent | Familiar | Lemon | 4 | Est-ce que ce citron est coupé ? | Is this lemon cut open? | yes |
| Referent | Familiar | Lemon | 4 | Est-ce que ce citron est tâché ? | Is this lemon stained? | no |
| Referent | Familiar | Mug | 1 | Est-ce que cette tasse est brillante ? | Is this mug shiny? | yes |
| Referent | Familiar | Mug | 1 | Est-ce que cette tasse est cassée ? | Is this mug broken? | no |
| Referent | Familiar | Mug | 2 | Est-ce que cette tasse est rouge ? | Is this mug red? | yes |
| Referent | Familiar | Mug | 2 | Est-ce que cette tasse est à l'envers ? | Is this mug upside down? | no |
| Referent | Familiar | Mug | 3 | Est-ce que la poignée de cette tasse est à droite ? | Is the mug handle on the right? | yes |
| Referent | Familiar | Mug | 3 | Est-ce que cette tasse est verte ? | Is this mug green? | no |
| Referent | Familiar | Mug | 4 | Est-ce que cette tasse est ronde ? | Is this mug round? | yes |
| Referent | Familiar | Mug | 4 | Est-ce que cette tasse a une queue ? | Does this mug have a stem? | no |
| Referent | Familiar | Strawberry | 1 | Est-ce que cette fraise a une queue ? | Does this strawberry have a stem? | yes |
| Referent | Familiar | Strawberry | 1 | Est-ce que cette fraise est verte ? | Is this strawberry green? | no |
| Referent | Familiar | Strawberry | 2 | Est-ce que cette fraise est rouge ? | Is this strawberry red? | yes |
| Referent | Familiar | Strawberry | 2 | Est-ce que cette fraise est coupée en morceaux ? | Is this strawberry cut into pieces? | no |
| Referent | Familiar | Strawberry | 3 | Est-ce qu'il y a deux fraises ? | Are there two strawberries? | yes |
| Referent | Familiar | Strawberry | 3 | Est-ce qu'il n'y a qu'une seule fraise ? | Is there only one strawberry? | no |
| Referent | Familiar | Strawberry | 4 | Est-ce que cette fraise est brillante ? | Is this strawberry shiny? | yes |
| Referent | Familiar | Strawberry | 4 | Est-ce que cette fraise est accrochée au fraisier ? | Is this strawberry attached to the plant? | no |
| Referent | Familiar | Sheep | 1 | Est-ce que ce mouton a de la laine ? | Does this sheep have wool? | yes |
| Referent | Familiar | Sheep | 1 | Est-ce que ce mouton est noir ? | Is this sheep black? | no |
| Referent | Familiar | Sheep | 2 | Est-ce que ce mouton a des oreilles ? | Does this sheep have ears? | yes |
| Referent | Familiar | Sheep | 2 | Est-ce que ce mouton a une queue ? | Does this sheep have a tail? | no |
| Referent | Familiar | Sheep | 3 | Est-ce que ce mouton est debout ? | Is this sheep standing? | yes |
| Referent | Familiar | Sheep | 3 | Est-ce que ce mouton a une corne ? | Does this sheep have a horn? | no |
| Referent | Familiar | Sheep | 4 | Est-ce que ce mouton a 4 pattes ? | Does this sheep have 4 legs? | yes |
| Referent | Familiar | Sheep | 4 | Est-ce que ce mouton est assis ? | Is this sheep sitting? | no |
| Referent | Familiar | Butterfly | 1 | Est-ce que ce papillon a des ailes ? | Does this butterfly have wings? | yes |
| Referent | Familiar | Butterfly | 1 | Est-ce que ce papillon a les ailes fermées ? | Does this butterfly have its wings closed? | no |
| Referent | Familiar | Butterfly | 2 | Est-ce que ce papillon a des points rouges sur les ailes ? | Does this butterfly have red spots on its wings? | yes |
| Referent | Familiar | Butterfly | 2 | Est-ce que ce papillon a une seule couleur ? | Is this butterfly a single color? | no |
| Referent | Familiar | Butterfly | 3 | Est-ce que ce papillon a plusieurs couleurs ? | Does this butterfly have multiple colors? | yes |
| Referent | Familiar | Butterfly | 3 | Est-ce que ce papillon est transparent ? | Is this butterfly transparent? | no |
| Referent | Familiar | Butterfly | 4 | Est-ce que ce papillon a des antennes ? | Does this butterfly have antennae? | yes |
| Referent | Familiar | Butterfly | 4 | Est-ce que ce papillon a une queue ? | Does this butterfly have a tail? | no |
| Referent | Familiar | Apple | 1 | Est-ce que cette pomme est ronde ? | Is this apple round? | yes |
| Referent | Familiar | Apple | 1 | Est-ce que cette pomme est verte ? | Is this apple green? | no |

| | | | | | | |
|---|---|---|---|---|---|---|
| Referent | Familiar | Apple | 2 | Est-ce que cette pomme a de la peau ? | Does this apple have skin? | yes |
| Referent | Familiar | Apple | 2 | Est-ce que cette pomme est coupée ? | Is this apple cut? | no |
| Referent | Familiar | Apple | 3 | Est-ce que cette pomme est rouge ? | Is this apple red? | yes |
| Referent | Familiar | Apple | 3 | Est-ce que cette pomme a des feuilles ? | Does this apple have leaves? | no |
| Referent | Familiar | Apple | 4 | Est-ce que cette pomme a une queue ? | Does this apple have a stem? | yes |
| Referent | Familiar | Apple | 4 | Est-ce que cette pomme est écrasée ? | Is this apple crushed? | no |
| Referent | Familiar | Banana | 1 | Est-ce que cette banane est jaune ? | Is this banana yellow? | yes |
| Referent | Familiar | Banana | 1 | Est-ce que cette banane est verte ? | Is this banana green? | no |
| Referent | Familiar | Banana | 2 | Est-ce que cette banane pointe vers le haut ? | Is this banana pointing upward? | yes |
| Referent | Familiar | Banana | 2 | Est-ce que cette banane pointe vers le bas ? | Is this banana pointing downward? | no |
| Referent | Familiar | Banana | 3 | Est-ce qu'il n'y a qu'une seule banane ? | Is there only one banana? | yes |
| Referent | Familiar | Banana | 3 | Est-ce qu'il y a plusieurs bananes ? | Are there multiple bananas? | no |
| Referent | Familiar | Banana | 4 | Est-ce que cette banane a de la peau ? | Does this banana have its peel? | yes |
| Referent | Familiar | Banana | 4 | Est-ce que cette banane est épluchée ? | Is this banana peeled? | no |
| Referent | Familiar | Turtle | 1 | Est-ce que cette tortue a des nageoires ? | Does this turtle have flippers? | yes |
| Referent | Familiar | Turtle | 1 | Est-ce que cette tortue a une queue ? | Does this turtle have a tail? | no |
| Referent | Familiar | Turtle | 2 | Est-ce que cette tortue a une carapace ? | Does this turtle have a shell? | yes |
| Referent | Familiar | Turtle | 2 | Est-ce que cette tortue est bleue ? | Is this turtle blue? | no |
| Referent | Familiar | Turtle | 3 | Est-ce que cette tortue a des écailles ? | Does this turtle have scales? | yes |
| Referent | Familiar | Turtle | 3 | Est-ce que cette tortue a une corne ? | Does this turtle have a horn? | no |
| Referent | Familiar | Turtle | 4 | Est-ce que cette tortue regarde vers la droite ? | Is this turtle looking to the right? | yes |
| Referent | Familiar | Turtle | 4 | Est-ce que cette tortue a des poils ? | Does this turtle have fur? | no |
| Even | Novel | Apoite | 1 | Est-ce que cet apoite a des pieds ? | Does this apoite have feet? | yes |
| Even | Novel | Apoite | 1 | Est-ce que cet apoite est vivant ? | Is this apoite alive? | no |
| Even | Novel | Apoite | 2 | Est-ce que cet apoite est en bois ? | Is this apoite made of wood? | yes |
| Even | Novel | Apoite | 2 | Est-ce que cet apoite roule ? | Does this apoite roll? | no |
| Even | Novel | Apoite | 3 | Est-ce que cet apoite est un objet ? | Is this apoite an object? | yes |
| Even | Novel | Apoite | 3 | Est-ce que cet apoite a des cornes ? | Does this apoite have horns? | no |
| Even | Novel | Apoite | 4 | Est-ce que cet apoite est marron ? | Is this apoite brown? | yes |
| Even | Novel | Apoite | 4 | Est-ce que cet apoite est rouge ? | Is this apoite red? | no |
| Even | Novel | Brime | 1 | Est-ce que cette brime est jaune ? | Is this brime yellow? | yes |
| Even | Novel | Brime | 1 | Est-ce que cette brime a des ailes ? | Does this brime have wings? | no |
| Even | Novel | Brime | 2 | Est-ce qu'il y a plusieurs brimes ? | Are there multiple brimes? | yes |
| Even | Novel | Brime | 2 | Est-ce que cette brime est un objet ? | Is this brime an object? | no |
| Even | Novel | Brime | 3 | Est-ce que cette brime a des épis ? | Does this brime have spikes? | yes |
| Even | Novel | Brime | 3 | Est-ce que cette brime peut courir ? | Can this brime run? | no |
| Even | Novel | Brime | 4 | Est-ce que cette brime est naturelle ? | Is this brime natural? | yes |
| Even | Novel | Brime | 4 | Est-ce que cette brime a un bec ? | Does this brime have a beak? | no |
| Even | Novel | Caralle | 1 | Est-ce que cette caralle a des feuilles ? | Does this caralle have leaves? | yes |
| Even | Novel | Caralle | 1 | Est-ce que cette caralle a des pattes ? | Does this caralle have legs? | no |
| Even | Novel | Caralle | 2 | Est-ce que cette caralle est un fruit ? | Is this caralle a fruit? | yes |
| Even | Novel | Caralle | 2 | Est-ce que cette caralle a de la fourrure ? | Does this caralle have fur? | no |
| Even | Novel | Caralle | 3 | Est-ce que cette caralle est rose ? | Is this caralle pink? | yes |
| Even | Novel | Caralle | 3 | Est-ce que cette caralle est jaune ? | Is this caralle yellow? | no |
| Even | Novel | Caralle | 4 | Est-ce que cette caralle a des noyaux ? | Does this caralle have seeds? | yes |
| Even | Novel | Caralle | 4 | Est-ce que cette caralle est fabriqué par l'homme ? | Is this caralle man-made? | no |
| Even | Novel | Civane | 1 | Est-ce que cette civane a des pattes ? | Does this civane have legs? | yes |
| Even | Novel | Civane | 1 | Est-ce que cette civane peut voler ? | Can this civane fly? | no |
| Even | Novel | Civane | 2 | Est-ce que cette civane est poilue ? | Is this civane furry? | yes |
| Even | Novel | Civane | 2 | Est-ce que cette civane est un objet ? | Is this civane an object? | no |
| Even | Novel | Civane | 3 | Est-ce que cette civane a des pinces ? | Does this civane have claws? | yes |
| Even | Novel | Civane | 3 | Est-ce que cette civane a un bec ? | Does this civane have a beak? | no |
| Even | Novel | Civane | 4 | Est-ce que cette civane est blanche ? | Is this civane white? | yes |
| Even | Novel | Civane | 4 | Est-ce que cette civane a des tâches ? | Does this civane have spots? | no |
| Even | Novel | Cratorre | 1 | Est-ce que ce cratorre a de grands yeux ? | Does this cratorre have large eyes? | yes |

| | | | | | | |
|---|---|---|---|---|---|---|
| Even | Novel | Cratorre | 1 | Est-ce que ce cratorre a des écailles ? | Does this cratorre have scales? | no |
| Even | Novel | Cratorre | 2 | Est-ce que ce cratorre est un animal ? | Is this cratorre an animal? | yes |
| Even | Novel | Cratorre | 2 | Est-ce que ce cratorre a des cornes ? | Does this cratorre have horns? | no |
| Even | Novel | Cratorre | 3 | Est-ce que ce cratorre a de la fourrure ? | Does this cratorre have fur? | yes |
| Even | Novel | Cratorre | 3 | Est-ce que ce cratorre est en bois ? | Is this cratorre made of wood? | no |
| Even | Novel | Cratorre | 4 | Est-ce que ce cratorre peut grimper aux arbres ? | Can this cratorre climb trees? | yes |
| Even | Novel | Cratorre | 4 | Est-ce que ce cratorre est rouge ? | Is this cratorre red? | no |
| Even | Novel | Guion | 1 | Est-ce que ce guion a des feuilles ? | Does this guion have leaves? | yes |
| Even | Novel | Guion | 1 | Est-ce que ce guion a des poils ? | Does this guion have fur? | no |
| Even | Novel | Guion | 2 | Est-ce que ce guion a une coque ? | Does this guion have a shell? | yes |
| Even | Novel | Guion | 2 | Est-ce que ce guion a des cornes ? | Does this guion have horns? | no |
| Even | Novel | Guion | 3 | Est-ce que ce guion est blanc à l'intérieur ? | Is this guion white on the inside? | yes |
| Even | Novel | Guion | 3 | Est-ce que ce guion est rose ? | Is this guion pink? | no |
| Even | Novel | Guion | 4 | Est-ce que ce guion est naturel ? | Is this guion natural? | yes |
| Even | Novel | Guion | 4 | Est-ce que ce guion peut courir ? | Can this guion run? | no |
| Even | Novel | Leipe | 1 | Est-ce que cette leipe vole ? | Does this leipe fly? | yes |
| Even | Novel | Leipe | 1 | Est-ce que cette leipe a des nageoires ? | Does this leipe have fins? | no |
| Even | Novel | Leipe | 2 | Est-ce que cette leipe a des pattes ? | Does this leipe have legs? | yes |
| Even | Novel | Leipe | 2 | Est-ce que cette leipe a des griffes ? | Does this leipe have claws? | no |
| Even | Novel | Leipe | 3 | Est-ce que cette leipe a un dard ? | Does this leipe have a stinger? | yes |
| Even | Novel | Leipe | 3 | Est-ce que cette leipe rampe ? | Does this leipe crawl? | no |
| Even | Novel | Leipe | 4 | Est-ce que cette leipe est bleue ? | Is this leipe blue? | yes |
| Even | Novel | Leipe | 4 | Est-ce que cette leipe a des écailles ? | Does this leipe have scales? | no |
| Even | Novel | Pesse | 1 | Est-ce que cette pesse a des pépins ? | Does this pesse have seeds? | yes |
| Even | Novel | Pesse | 1 | Est-ce que cette pesse a des ailes ? | Does this pesse have wings? | no |
| Even | Novel | Pesse | 2 | Est-ce que cette pesse a une coque verte ? | Does this pesse have a green shell? | yes |
| Even | Novel | Pesse | 2 | Est-ce que cette pesse a des poils ? | Does this pesse have fur? | no |
| Even | Novel | Pesse | 3 | Est-ce que cette pesse est blanche à l'intérieur ? | Is this pesse white on the inside? | yes |
| Even | Novel | Pesse | 3 | Est-ce que cette pesse est un animal ? | Is this pesse an animal? | no |
| Even | Novel | Pesse | 4 | Est-ce que cette pesse est naturelle ? | Is this pesse natural? | yes |
| Even | Novel | Pesse | 4 | Est-ce que cette pesse est rouge ? | Is this pesse red? | no |
| Even | Novel | Phot | 1 | Est-ce que ce phot a des yeux ? | Does this phot have eyes? | yes |
| Even | Novel | Phot | 1 | Est-ce que ce phot a des poils ? | Does this phot have fur? | no |
| Even | Novel | Phot | 2 | Est-ce que ce phot nage ? | Does this phot swim? | yes |
| Even | Novel | Phot | 2 | Est-ce que ce phot peut voler ? | Can this phot fly? | no |
| Even | Novel | Phot | 3 | Est-ce que ce phot est bleu ? | Is this phot blue? | yes |
| Even | Novel | Phot | 3 | Est-ce que ce phot est jaune ? | Is this phot yellow? | no |
| Even | Novel | Phot | 4 | Est-ce que ce phot a deux nageoires ? | Does this phot have two fins? | yes |
| Even | Novel | Phot | 4 | Est-ce que ce phot a des pattes ? | Does this phot have legs? | no |
| Even | Novel | Poimon | 1 | Est-ce que ce poimon a un manche ? | Does this poimon have a handle? | yes |
| Even | Novel | Poimon | 1 | Est-ce que ce poimon est noir ? | Is this poimon black? | no |
| Even | Novel | Poimon | 2 | Est-ce que ce poimon a une boule au bout ? | Does this poimon have a ball at the end? | yes |
| Even | Novel | Poimon | 2 | Est-ce que ce poimon a des ailes ? | Does this poimon have wings? | no |
| Even | Novel | Poimon | 3 | Est-ce que ce poimon est brillant ? | Is this poimon shiny? | yes |
| Even | Novel | Poimon | 3 | Est-ce que ce poimon a des pattes ? | Does this poimon have legs? | no |
| Even | Novel | Poimon | 4 | Est-ce que ce poimon est un objet ? | Is this poimon an object? | yes |
| Even | Novel | Poimon | 4 | Est-ce que ce poimon a des yeux ? | Does this poimon have eyes? | no |
| Even | Novel | Setrin | 1 | Est-ce que ce setrin a des pics ? | Does this setrin have spikes? | yes |
| Even | Novel | Setrin | 1 | Est-ce que ce setrin a des pattes ? | Does this setrin have legs? | no |
| Even | Novel | Setrin | 2 | Est-ce que ce setrin rampe ? | Does this setrin crawl? | yes |
| Even | Novel | Setrin | 2 | Est-ce que ce setrin est bleu ? | Is this setrin blue? | no |
| Even | Novel | Setrin | 3 | Est-ce que ce setrin est rouge ? | Is this setrin red? | yes |
| Even | Novel | Setrin | 3 | Est-ce que ce setrin a des ailes ? | Does this setrin have wings? | no |
| Even | Novel | Setrin | 4 | Est-ce que la tête du setrin est face à vous ? | Is this setrin's head facing you? | yes |
| Even | Novel | Setrin | 4 | Est-ce que ce setrin est un objet ? | Is this setrin an object? | no |
| Even | Novel | Tosse | 1 | Est-ce que ce tosse a des cornes ? | Does this tosse have horns? | yes |
| Even | Novel | Tosse | 1 | Est-ce que ce tosse est vert ? | Is this tosse green? | no |
| Even | Novel | Tosse | 2 | Est-ce que les cornes du tosse sont-elles pointées vers le haut ? | Are this tosse's horns pointing upward? | yes |
| Even | Novel | Tosse | 2 | Est-ce que ce tosse a une queue ? | Does this tosse have a tail? | no |
| Even | Novel | Tosse | 3 | Est-ce que ce tosse a des poils ? | Does this tosse have fur? | yes |
| Even | Novel | Tosse | 3 | Est-ce que ce tosse a des plumes ? | Does this tosse have feathers? | no |
| Even | Novel | Tosse | 4 | Est-ce que ce tosse a des pattes ? | Does this tosse have legs? | yes |
| Even | Novel | Tosse | 4 | Est-ce que ce tosse a des rayures ? | Does this tosse have stripes? | no |
| Odd | Novel | Bardon | 1 | Est-ce que ce bardon a des pattes ? | Does this bardon have legs? | yes |

| | | | | | | |
|---|---|---|---|---|---|---|
| Odd | Novel | Bardon | 1 | Est-ce que ce bardon a des écailles ? | Does this bardon have scales? | no |
| Odd | Novel | Bardon | 2 | Est-ce que les oreilles du bardon pointent vers le haut ? | Are this bardon's ears pointing upward? | yes |
| Odd | Novel | Bardon | 2 | Est-ce que ce bardon est noir ? | Is this bardon black? | no |
| Odd | Novel | Bardon | 3 | Est-ce que ce bardon a un museau ? | Does this bardon have a snout? | yes |
| Odd | Novel | Bardon | 3 | Est-ce que ce bardon regarde vers la gauche ? | Is this bardon looking to the left? | no |
| Odd | Novel | Bardon | 4 | Est-ce que ce bardon a des poils clairs ? | Does this bardon have light-colored fur? | yes |
| Odd | Novel | Bardon | 4 | Est-ce que ce bardon a des ailes ? | Does this bardon have wings? | no |
| Odd | Novel | Edoime | 1 | Est-ce que l'edoime a des poils ? | Does this edoime have fur? | yes |
| Odd | Novel | Edoime | 1 | Est-ce que l'edoime a des ailes ? | Does this edoime have wings? | no |
| Odd | Novel | Edoime | 2 | Est-ce que l'edoime a des pattes ? | Does this edoime have legs? | yes |
| Odd | Novel | Edoime | 2 | Est-ce que l'edoime est bleu ? | Is this edoime blue? | no |
| Odd | Novel | Edoime | 3 | Est-ce que l'edoime regarde vers la gauche ? | Is this edoime looking to the left? | yes |
| Odd | Novel | Edoime | 3 | Est-ce que l'edoime est en bois ? | Is this edoime made of wood? | no |
| Odd | Novel | Edoime | 4 | Est-ce que l'edoime est noir et blanc ? | Is this edoime black and white? | yes |
| Odd | Novel | Edoime | 4 | Est-ce que l'edoime a une corne ? | Does this edoime have a horn? | no |
| Odd | Novel | Crute | 1 | Est-ce que cette crute est blanche ? | Is this crute white? | yes |
| Odd | Novel | Crute | 1 | Est-ce que cette crute a des pattes ? | Does this crute have legs? | no |
| Odd | Novel | Crute | 2 | Est-ce que cette crute a une tige ? | Does this crute have a stem? | yes |
| Odd | Novel | Crute | 2 | Est-ce que cette crute est coupée en deux ? | Is this crute cut in half? | no |
| Odd | Novel | Crute | 3 | Est-ce que cette crute a des tâches ? | Does this crute have spots? | yes |
| Odd | Novel | Crute | 3 | Est-ce que cette crute est verte ? | Is this crute green? | no |
| Odd | Novel | Crute | 4 | Est-ce que cette crute est ovale ? | Is this crute oval-shaped? | yes |
| Odd | Novel | Crute | 4 | Est-ce que cette crute a des plumes ? | Does this crute have feathers? | no |
| Odd | Novel | Toupe | 1 | Est-ce que cette toupe est coupée ? | Is this toupe cut? | yes |
| Odd | Novel | Toupe | 1 | Est-ce que cette toupe est rouge ? | Is this toupe red? | no |
| Odd | Novel | Toupe | 2 | Est-ce qu'il y a plusieurs toupes ? | Are there multiple toupes? | yes |
| Odd | Novel | Toupe | 2 | Est-ce que cette toupe a des plumes ? | Does this toupe have feathers? | no |
| Odd | Novel | Toupe | 3 | Est-ce que cette toupe a une peau ? | Does this toupe have a peel? | yes |
| Odd | Novel | Toupe | 3 | Est-ce que cette toupe est carrée ? | Is this toupe square? | no |
| Odd | Novel | Toupe | 4 | Est-ce que cette toupe a des feuilles ? | Does this toupe have leaves? | yes |
| Odd | Novel | Toupe | 4 | Est-ce que cette toupe est en bois ? | Is this toupe made of wood? | no |
| Odd | Novel | Loppe | 1 | Est-ce que ce loppe a une manivelle ? | Does this loppe have a crank? | yes |
| Odd | Novel | Loppe | 1 | Est-ce que ce loppe est en bois ? | Is this loppe made of wood? | no |
| Odd | Novel | Loppe | 2 | Est-ce que ce loppe a un pied ? | Does this loppe have a base? | yes |
| Odd | Novel | Loppe | 2 | Est-ce que ce loppe a des ailes ? | Does this loppe have wings? | no |
| Odd | Novel | Loppe | 3 | Est-ce que ce loppe est rouillé ? | Is this loppe rusty? | yes |
| Odd | Novel | Loppe | 3 | Est-ce que ce loppe est bleu ? | Is this loppe blue? | no |
| Odd | Novel | Loppe | 4 | Est-ce que ce loppe est un objet ? | Is this loppe an object? | yes |
| Odd | Novel | Loppe | 4 | Est-ce que ce loppe a des poils ? | Does this loppe have fur? | no |
| Odd | Novel | Madrin | 1 | Est-ce que ce madrin a un bec ? | Does this madrin have a beak? | yes |
| Odd | Novel | Madrin | 1 | Est-ce que ce madrin est bleu ? | Is this madrin blue? | no |
| Odd | Novel | Madrin | 2 | Est-ce que ce madrin a des ailes ? | Does this madrin have wings? | yes |
| Odd | Novel | Madrin | 2 | Est-ce que ce madrin regarde vers la gauche ? | Is this madrin looking to the left? | no |
| Odd | Novel | Madrin | 3 | Est-ce que ce madrin a des plumes ? | Does this madrin have feathers? | yes |
| Odd | Novel | Madrin | 3 | Est-ce que ce madrin a des écailles ? | Does this madrin have scales? | no |
| Odd | Novel | Madrin | 4 | Est-ce que ce madrin a une queue ? | Does this madrin have a tail? | yes |
| Odd | Novel | Madrin | 4 | Est-ce que ce madrin est en bois ? | Is this madrin made of wood? | no |
| Odd | Novel | Musse | 1 | Est-ce que cette musse est blanche à l'intérieur ? | Is this musse white on the inside? | yes |
| Odd | Novel | Musse | 1 | Est-ce que cette musse a des ailes ? | Does this musse have wings? | no |
| Odd | Novel | Musse | 2 | Est-ce que cette musse a une peau rose ? | Does this musse have pink skin? | yes |
| Odd | Novel | Musse | 2 | Est-ce que cette musse est noire ? | Is this musse black? | no |
| Odd | Novel | Musse | 3 | Est-ce qu'il y a plusieurs musses ? | Are there multiple musses? | yes |
| Odd | Novel | Musse | 3 | Est-ce que cette musse a des épines ? | Does this musse have thorns? | no |
| Odd | Novel | Musse | 4 | Est-ce que cette musse a des feuilles ? | Does this musse have leaves? | yes |
| Odd | Novel | Musse | 4 | Est-ce que cette musse a des écailles ? | Does this musse have scales? | no |
| Odd | Novel | Nuraque | 1 | Est-ce que ce nuraque a une peau orange ? | Does this nuraque have orange skin? | yes |
| Odd | Novel | Nuraque | 1 | Est-ce que ce nuraque est violet ? | Is this nuraque purple? | no |
| Odd | Novel | Nuraque | 2 | Est-ce que ce nuraque est vert à l'intérieur ? | Is this nuraque green on the inside? | yes |
| Odd | Novel | Nuraque | 2 | Est-ce que ce nuraque a des ailes ? | Does this nuraque have wings? | no |
| Odd | Novel | Nuraque | 3 | Est-ce que ce nuraque a des pics ? | Does this nuraque have spikes? | yes |
| Odd | Novel | Nuraque | 3 | Est-ce que ce nuraque a des cornes ? | Does this nuraque have horns? | no |

| | | | | | | |
|---|---|---|---|---|---|---|
| Odd | Novel | Nuraque | 4 | Est-ce que ce nuraque est un fruit ? | Is this nuraque a fruit? | yes |
| Odd | Novel | Nuraque | 4 | Est-ce que ce nuraque est en bois ? | Is this nuraque made of wood? | no |
| Odd | Novel | Plaime | 1 | Est-ce que cette plaime a une queue ? | Does this plaime have a tail? | yes |
| Odd | Novel | Plaime | 1 | Est-ce que cette plaime est verte ? | Is this plaime green? | no |
| Odd | Novel | Plaime | 2 | Est-ce que cette plaime est bleue ? | Is this plaime blue? | yes |
| Odd | Novel | Plaime | 2 | Est-ce que cette plaime a des plumes ? | Does this plaime have feathers? | no |
| Odd | Novel | Plaime | 3 | Est-ce que cette plaime a un long corps ? | Does this plaime have a long body? | yes |
| Odd | Novel | Plaime | 3 | Est-ce que cette plaime a des cornes ? | Does this plaime have horns? | no |
| Odd | Novel | Plaime | 4 | Est-ce que cette plaime a des bras ? | Does this plaime have arms? | yes |
| Odd | Novel | Plaime | 4 | Est-ce que cette plaime a des poils ? | Does this plaime have fur? | no |
| Odd | Novel | Possecale | 1 | Est-ce que ce possecale a un corps en forme de tube ? | Does this possecale have a tube-shaped body? | yes |
| Odd | Novel | Possecale | 1 | Est-ce que ce possecale est vert ? | Is this possecale green? | no |
| Odd | Novel | Possecale | 2 | Est-ce que ce possecale a des épines ? | Does this possecale have spines? | yes |
| Odd | Novel | Possecale | 2 | Est-ce que ce possecale a des yeux ? | Does this possecale have eyes? | no |
| Odd | Novel | Possecale | 3 | Est-ce que ce possecale a des rainures ? | Does this possecale have grooves? | yes |
| Odd | Novel | Possecale | 3 | Est-ce que ce possecale est en bois ? | Is this possecale made of wood? | no |
| Odd | Novel | Possecale | 4 | Est-ce que ce possecale est orange à la base ? | Is this possecale orange at the base? | yes |
| Odd | Novel | Possecale | 4 | Est-ce que ce possecale a de la fourrure ? | Does this possecale have fur? | no |
| Odd | Novel | Rapare | 1 | Est-ce que ce rapare a les yeux rouges ? | Does this rapare have red eyes? | yes |
| Odd | Novel | Rapare | 1 | Est-ce que ce rapare a des ailes ? | Does this rapare have wings? | no |
| Odd | Novel | Rapare | 2 | Est-ce que ce rapare a des antennes ? | Does this rapare have antennae? | yes |
| Odd | Novel | Rapare | 2 | Est-ce que ce rapare est jaune ? | Is this rapare yellow? | no |
| Odd | Novel | Rapare | 3 | Est-ce que ce rapare est bleu ? | Is this rapare blue? | yes |
| Odd | Novel | Rapare | 3 | Est-ce que ce rapare peut marcher ? | Can this rapare walk? | no |
| Odd | Novel | Rapare | 4 | Est-ce que ce rapare a des moustaches ? | Does this rapare have whiskers? | yes |
| Odd | Novel | Rapare | 4 | Est-ce que ce rapare est un fruit ? | Is this rapare a fruit? | no |
| Odd | Novel | Thiou | 1 | Es-ce que ce thiou est doré ? | Is this thiou golden? | yes |
| Odd | Novel | Thiou | 1 | Est-ce que ce thiou a des jambes ? | Does this thiou have legs? | no |
| Odd | Novel | Thiou | 2 | Est-ce que ce thiou a une chaîne ? | Does this thiou have a chain? | yes |
| Odd | Novel | Thiou | 2 | Est-ce que ce thiou est noir ? | Is this thiou black? | no |
| Odd | Novel | Thiou | 3 | Est-ce que la partie centrale du thiou est ronde ? | Is the central part of this thiou round? | yes |
| Odd | Novel | Thiou | 3 | Est-ce que ce thiou a des poils ? | Does this thiou have fur? | no |
| Odd | Novel | Thiou | 4 | Est-ce que ce thiou est en métal ? | Is this thiou made of metal? | yes |
| Odd | Novel | Thiou | 4 | Est-ce que ce thiou est triangulaire ? | Is this thiou triangular? | no |

**Table S2. Associations between neuropsychological measures and memory task performance.**

Nominal correlations between neuropsychological measures and memory task performance indices in older (OA) and younger adults (YA). Only associations reaching $p < .05$ are reported. Memory task performance indices include accuracy measures from the DIILT learning phase (AUC for familiar items, AUC for novel items, CV for familiar items, CV for novel items, accuracy gap for Familiar-Novel items at Block 1 and Block 4) and recognition accuracy (immediate and delayed recognitions for implicit and explicit tasks). Correlations were computed using Pearson or Spearman methods depending on normality assumptions. *p*FDR values were adjusted using the Benjamini-Hochberg procedure within each group.

At the nominal level, in younger adults, higher accuracy for novel items during learning was positively associated with working memory (digit span backward: $r = .46$, $p = .011$; N-Back x Familiar-Novel accuracy gap at Block 4: $r = -.47$, $p = .007$) and episodic memory (CVLT cued immediate: $r = .39$, $p = .034$; CVLT cued delayed: $r = .38$, $p = .037$). In older adults, better immediate DIILT recognition was positively associated with processing speed ($r = .40$, $p = .020$). Additionally, older adults with better episodic memory showed a smaller Familiar-Novel accuracy gap at Block 1 (CVLT recognition: $r = -.39$, $p = .025$; CVLT delayed free recall: $r = -.35$, $p = .042$). Higher depressive symptoms were associated with lower recognition accuracy for associations learned implicitly (GDS x DIILT immediate recognition: $r = -.46$, $p = .007$) and explicitly (GDS x Explicit delayed recognition: $r = -.35$, $p = .039$) and a smaller Familiar-Novel accuracy gap at the end of learning (HADS Depression x Familiar-Novel gap at Block 4: $r = -.42$, $p = .014$). None of these associations survived FDR correction (all *p*FDR ≥ .466).

| Group | Memory task performance | Neuropsychological measure | n | *r* | *p* | *p*FDR |
|---|---|---|---|---|---|---|
| OA | DIILT immediate recognition accuracy | GDS | 33 | -.46 | .007** | .604 |
| OA | DIILT immediate recognition accuracy | Processing speed | 34 | .40 | .020* | .809 |
| OA | DIILT immediate recognition accuracy | ASQ Internal Orientation | 33 | -.36 | .040* | .830 |
| OA | Explicit encoding - immediate recognition accuracy | Age | 35 | -.34 | .049* | .830 |
| OA | Explicit encoding - delayed recognition accuracy | GDS | 34 | -.35 | .039* | .830 |
| OA | Explicit encoding - delayed recognition accuracy | BIS-11 motor impulsivity | 33 | .35 | .044* | .830 |
| OA | DIILT - AUC for familiar items | Age | 34 | -.52 | .002** | .466 |
| OA | DIILT - AUC for familiar items | HADS Depression | 33 | -.40 | .022* | .809 |
| OA | DIILT - CV for familiar items | BIS-11 Planning difficulties | 33 | -.48 | .004** | .579 |
| OA | DIILT - CV for familiar items | BIS-11 total score | 33 | -.39 | .025* | .809 |
| OA | DIILT - delta Familiar-Novel at Block 1 | CVLT recognition | 33 | -.39 | .025* | .809 |
| OA | DIILT - delta Familiar-Novel at Block 1 | BIS-11 motor impulsivity | 32 | -.38 | .030* | .830 |
| OA | DIILT - delta Familiar-Novel at Block 1 | CVLT free recall (delayed) | 34 | -.35 | .042* | .830 |
| OA | DIILT - delta Familiar-Novel at Block 1 | BIS-11 cognitive impulsivity | 33 | -.35 | .045* | .830 |

| | | | | | | |
|---|---|---|---|---|---|---|
| OA | DIILT - delta Familiar-Novel at Block 4 | HADS Depression | 34 | -.42 | .014* | .809 |
| YA | DIILT immediate recognition accuracy | Socioeconomic Status | 30 | .38 | .038* | .882 |
| YA | DIILT - AUC for familiar items | Digit span (backward) | 30 | .37 | .047* | .882 |
| YA | DIILT - AUC for novel items | Digit span (backward) | 30 | .46 | .011* | .882 |
| YA | DIILT - AUC for novel items | CVLT cued (immediate) | 30 | .39 | .034* | .882 |
| YA | DIILT - AUC for novel items | CVLT cued (delayed) | 30 | .38 | .037* | .882 |
| YA | DIILT - AUC for novel items | GDS | 30 | -.37 | .044* | .882 |
| YA | DIILT - CV for familiar items | BIS-11 total score | 31 | .45 | .011* | .882 |
| YA | DIILT - CV for novel items | Digit span (backward) | 31 | -.37 | .040* | .882 |
| YA | DIILT - CV for novel items | CVLT recognition | 31 | -.36 | .045* | .882 |
| YA | DIILT - delta Familiar-Novel at Block 1 | Digit span (backward) | 29 | -.41 | .027* | .882 |
| YA | DIILT - delta Familiar-Novel at Block 4 | N-Back | 31 | -.47 | .007** | .882 |
| YA | DIILT - delta Familiar-Novel at Block 4 | BIS-11 total score | 31 | -.39 | .029* | .882 |
| YA | DIILT - delta Familiar-Novel at Block 4 | BIS-11 cognitive impulsivity | 31 | -.39 | .029* | .882 |

**Note.** *r* = correlation coefficient; *p* = nominal *p*-value; *p*FDR = FDR-corrected *p*-value. AUC = Area Under the Curve; CV = Coefficient of Variation; YA = Younger Adults; OA = Older Adults. Significance levels are indicated as follows: $p < .05$*, $p < .01$**, $p < .001$***.

**Figure S1. Associations between cognitive abilities and implicit learning performance.**

(A) In younger adults, working memory scores (Digit Span Backward) are positively associated with accuracy for novel associations across learning, while no such association is observed in older adults.

(B) In older adults, better episodic memory (CVLT delayed free recall) is associated with a smaller Familiar-Novel accuracy gap at Block 1, suggesting that those with greater episodic memory resources were less reliant on familiar associations from the outset of learning. No such association is observed in younger adults.

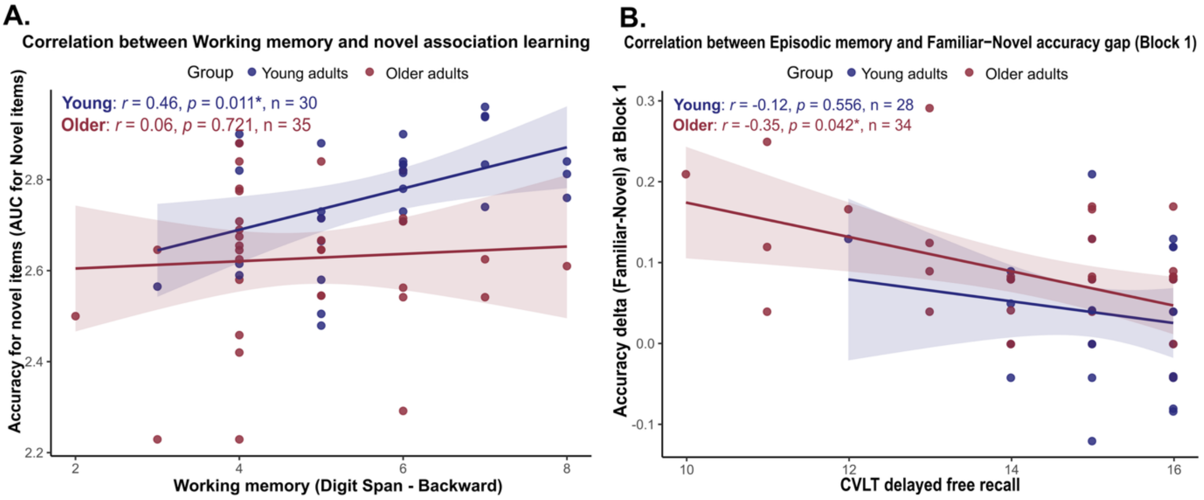


**Note**. *r* and *p* values correspond to Spearman or Pearson correlations depending on normality. All *p*s are nominal and did not survive FDR correction. Dark blue = younger adults; burgundy = older adults. Significance indicators are indicated as follows: $p < .05^*$, $p < .01^{**}$, $p < .001^{***}$.
